# Comprehensive Multi-Omic Analysis Reveals Distinct Molecular Features in Early and Advanced Stages of Hepatocellular Carcinoma

**DOI:** 10.1101/2023.12.17.570960

**Authors:** Mingzhu Fan, Jin Hu, Xiaoyan Xu, Jia Chen, Wenwen Zhang, Xiaoping Zheng, Jinheng Pan, Wei Xu, Shan Feng

**Affiliations:** Key Laboratory of Structural Biology of Zhejiang Province, Westlake University, Hangzhou 310024, Zhejiang, China; Mass Spectrometry & Metabolomics Core Facility, The Biomedical Research Core Facility, Westlake University, Hangzhou 310024, Zhejiang, China; College of Basic Medical Science, Zhejiang Chinese Medical University, Hangzhou 310053, Zhejiang, China; Key Laboratory of Chinese Medicine Rheumatology of Zhejiang Province, Zhejiang Chinese Medical University, Hangzhou 310053, Zhejiang, China; Hangzhou Tongchuang Medical laboratory, Shulan Health Group, Hangzhou 310015, Zhejiang, China; Pathology Department, Shulan (Hangzhou) Hospital, Hangzhou 311112, Zhejiang, China

**Author notes:** These authors contribute equally to this work (Mingzhu Fan, Jin Hu).

**Keywords:** Hepatocellular Carcinoma, Proteomics and Phosphoproteomics, Metabolomics and Lipidomics, Multi-omic analysis and integration

## Abstract

Hepatocellular Carcinoma (HCC) is a serious primary solid tumor that is prevalent worldwide. Due to its high mortality rate, it is crucial to explore both early diagnosis and advanced treatment for HCC. In recent years, multi-omic approaches have emerged as promising tools to identify biomarkers and investigate molecular mechanisms of biological processes and diseases. In this study, we performed proteomics, phosphoproteomics, metabolomics, and lipidomics to reveal the molecular features of early- and advanced-stage HCC. The data obtained from these omics were analyzed separately and then integrated to provide a comprehensive understanding of the disease. The multi-omic results unveiled intricate biological pathways and interaction networks underlying the initiation and progression of HCC. Moreover, we proposed specific potential biomarker panels for both early- and advanced-stage HCC by overlapping our data with CPTAC database, and deduced novel insights and mechanisms related to HCC origination and development.

## 1. Introduction

Hepatocellular Carcinoma (HCC) is the most common form of liver cancer and originates in hepatocytes. It is often associated with various liver diseases, such as cirrhosis or chronic hepatitis B or C infection [1-3]. HCC is now one of the most lethal cancer-induced diseases worldwide [4-5], and patients in developing countries have an average survival time of only a few months after clinical diagnosis via traditional medical imaging strategies, although there has been significant improvement in HCC survival rates in recent years [6]. Due to its heterogeneity, complexity, and malignancy, both early diagnosis and intervention for HCC remain challenging. Although molecular characterization of HCC tissue is increasingly important, alpha-fetoprotein (AFP), a glycoprotein produced by the fetal liver, is the only commonly used serum biomarker for diagnosis [7-9]. In contrast to diagnosis, the intervention methods for HCC often depend on the disease stage, which is classified as stages I to IV using the TNM (Tumor, Nodule & Metastasis) cancer staging system [4, 9-10]. The cancer stage is a crucial parameter used to describe the tumor’s growth and spreading.

Surgical resection and liver transplantation are commonly used to treat early-stage HCC, while radio ablation and systemic therapy are frequently used to treat advanced-stage HCC [3-4, 11-14]. For example, Sorafenib, the first multikinase inhibitor agent approved by the Food and Drug Administration (FDA), has significantly improved the survival of HCC patients [14-15]. However, its efficacy is limited by side effects, low response rates, and the development of drug resistance [15-16]. Although many other targeted and combinational therapies are in rapid development and Phase III clinical trials [17-18], a better understanding of the molecular mechanisms underlying HCC origin and progression is still needed to guide interventions.

Multi-omics integration is an emerging approach that combines high-throughput data from genomics, transcriptomics, proteomics, metabolomics, and lipidomics to provide a comprehensive understanding of HCC. Genomic and transcriptomic studies, which are based on whole-genome sequencing and RNA-seq or microarray assays, have identified numerous genetic alterations and dysregulated gene expression patterns associated with HCC [19-20]. Mass spectrometry-based proteomics can identify differentially expressed proteins in HCC-related cells or tissues compared to normal cells or tissues. Early proteomic work mostly focused on identifying prognostic biomarkers in serum or tumor tissue samples [21], while recent research has employed proteomics, phosphoproteomics, and AI-based data interpretation to uncover more mechanisms underlying HCC development and progression. This has led to the discovery of novel therapeutic targets, such as SOAT1, Src family kinase, PCNA, CDK1, FGFR4, MFAP5, etc [22-27].

Proteogenomic characterization is a multi-omics strategy that integrates genomic, transcriptomic, and proteomic data. It has emerged as a valuable approach in cancer research, providing biological and diagnostic insights [28]. For example, Gao et al. performed a proteogenomic study using 159 paired HBV-related HCC tissues and identified PYCR2 and ADH1A as two prognostic biomarkers. Similarly, Ng et al. used proteogenomic analysis to reveal dysregulation of Wnt-β-catenin, AKT/mTOR, and Notch pathways in HCC [29-30]. In addition to genomics, transcriptomics and proteomics, metabolomics and lipidomics are alternative tools for investigating biomarkers and therapeutic targets of HCC at the level of the final products of genes and proteins.

Compared to lipidomics, the metabolomic approach primarily focuses on hydrophilic metabolites, particularly those involved in central metabolism pathways. In addition to the well-known Warburg effect, which is a general characteristic of cancer cell metabolism, HCC exhibits a distinct metabolite profile compared to other cancers. Previous studies have shown that metabolic molecules involved in glycolysis, the TCA cycle, amino acid metabolism, and bile acid metabolism are disrupted in association with HCC [31-33]. Lipidomics is a complementary approach to metabolomics that primarily profiles endogenous hydrophobic compounds. Numerous studies have shown that alterations in phospholipid, sphingolipid, β-oxidation, and ketone body biosynthesis in HCC cells or tissues contribute to the initiation and development of the disease [34-35]. Recently, more sets of combinational biomarkers for premalignant liver disease have been identified and summarized using metabolomic and lipidomic approaches [36].

Numerous studies have been conducted to identify specific patterns associated with hepatocellular carcinoma (HCC) disease using single or multiple omics approaches. However, most studies have focused on early diagnosis and treatment, and research on advanced-stage HCC remains insufficient. To address this gap, we employed a multi-omics approach, including quantitative proteomics, phosphoproteomics, metabolomics, and lipidomics, to investigate differential patterns between early- and advanced-stage HCC tumors. We compared our results with CPTAC database and integrated the multi-omics data using QIAGEN Ingenuity Pathway Analysis (IPA), revealing the complexity of the pathway network during tumor progression. These findings not only provide valuable insights into liver cancer biology but also demonstrate the feasibility of integrating multi-dimensional information from macro biomolecules, protein translational modification, and small compounds.

## 2. Materials and Methods

### 2.1. Clinical sample collection and study cohort

All patients involved in this study provided informed consent for biobank recruitment at Shulan (Hangzhou) Hospital. Both tumor and adjacent non-tumor liver tissue samples were procured from hepatocellular carcinoma (HCC) patients during surgical procedures. These samples were carefully sectioned into pieces ranging from 0.3 to 0.8 cm3 using a scalpel. Following extraction, the fresh tissue samples were promptly frozen and preserved in liquid nitrogen prior to sample processing. Ethical clearance was granted by the Research Ethics Committee at Shulan (Hangzhou) Hospital.

The diagnosis of HCC was confirmed either through histopathological examination or by adhering to established imaging criteria. The clinical stage of the disease was subsequently ascertained based on the TNM cancer staging system. The study encompassed fourteen pairs of HCC and corresponding non-tumor liver tissue samples. Half of these pairs were derived from patients diagonosed with early-stage (TNM stage I, characterized by a solitary tumor of ≤ 2 cm with no vascular invasion), while the remining pairs were from patients with advanced-stage (TNM stage III & IV, marked by a solitary tumor of ≥ 2 cm or multiple tumors, vascular invasion, and in certain instances, distant metastasis). The statistical characteristics of patients with early-stage and advanced-stage HCC are depicted in Table 1.

**Tables 1.** Characteristics of the Cohorts.

|  |  | Early-stage | Advanced-stage |
| --- | --- | --- | --- |
| Age | Mean±SD | 60.3±11.9 | 46.4±13.5 |
| Gender | Male% | 57.1 | 42.8 |
| Smoking | Yes% | 28.6 | 28.6 |
|  | No% | 71.4 | 71.4 |
| Alcohol | Yes% | 0 | 28.6 |
|  | No% | 100 | 71.4 |
| Cirrhosis | Yes% | 14.3 | 14.3 |
| Cancer grade | Low% | 14.3 | 57.1 |
|  | Medium% | 85.7 | 42.9 |
| Tumor size | Mean±SD | 110.5±146.2 cm<sup>3</sup> | 684.1±582.9 cm<sup>3</sup> |
| Invasion | Yes% | 0 | 71 |
| Metastasis | Yes% | 0 | 57.1 |

All tissue samples were ground and homogenized using a cryogenic grinder (SPEX 6875 Freezer/Mill) and divided into four parts by weight: 3 mg for proteomic experiments, 80 mg for phosphoproteomic experiments, 50 mg for metabolite extraction, and 50 mg for lipid extraction.

### 2.2. Reagents, protein, metabolites, lipid extraction and phosphopeptide enrichment

All the detailed information were provided in supplementary part. Briefly, all the main reagents used in this study were from Sigma, Promega, Thermo Fisher, etc. Protein, metabolite and lipid were exacted by RIPA buffer, 80% methanol and Methyl tert-butyl ether (MTBE) from the ground tissue, respectively. Protein was digested by trypsin overnight and further labeled by TMT regent for proteomics, phosphopetide was enriched by IMAC Fe-NTA and TiO_2_.

### 2.3. LC-MS/MS methods

The TMT-labeled peptides were separated using a Thermo Vanquish Neo integrated nano-HPLC system directly interfaced with a Thermo Exploris 480 mass spectrometer equipped with FAIMS Pro. The enriched phosphopeptides were analyzed using a Bruker Nano Elute HPLC system directly interfaced with a Bruker timsTOF Pro 2 mass spectrometer. Both metabolomic and lipidomic analyses were performed using an Agilent 1290 Infinity HPLC system coupled to an Agilent 6545 Q-tof mass spectrometer with an electrospray ion source operating in both positive and negative ion modes. The detailed information can be found in supplementary part.

### 2.4. Bioinformatic analysis

Proteomic data were analyzed using Gene Ontology (GO) and Kyoto Encyclopedia of Genes and Genomes (KEGG) assays. GO enrichment analysis was conducted on the website (http://geneontology.org/) by importing the gene IDs of up-or down-regulated proteins [37, 38], and a cut-off less than 0.05 was used for GO category analysis after multiple test correction. KEGG pathway enrichment was performed on the website (http://www.genome.jp/kegg/kaas/) by importing the gene IDs of up- or down-regulated proteins with a 0.01 cut-off for p values [39].

A kinase-substrate relation was established using the KSEA app (casecpb.shinyapps.io/ksea/) based on phosphoproteomic data [40]. Only significantly changed (p<0.05) phosphopeptides were used for the assay, and the filtered output contained kinase predictions with a z-score of ≥2.0.

Survival analysis was performed on the website (http://gepia2.cancer-pku.cn/#survival) based on a multi-gene signature and plot a Kaplan-Meier curve.

To perform multi-omic integrative analysis, Ingenuity Pathway Analysis (IPA) was used. Significant up- or down-regulated proteins, metabolites, and lipids were combined into a single sheet, which was uploaded to IPA for core analysis. The output parameters of pathway enrichment and network building were set to default. The data of dysregulated phosphopeptides were uploaded separately for phosphorylation analysis. Comparison analysis was used to summarize the enriched pathways from the protein, metabolite, lipid, and phosphorylation assays for both early and advanced-stage HCC.

### 2.5. Statistics

Data were presented following the analysis of variance (ANOVA). To calculate the statistical significance between groups, Student’s t-test, assuming two-tailed distributions, or the nonparametric Mann-Whitney test was used, without assuming that the variance was similar between the groups being statistically compared. A p-value of <0.05 was considered statistically significant.

## 3. Results

LC-MS/MS data of TMT-labeled quantitative proteomics, phosphoproteomics, metabolomics, and lipidomics for the same HCC tumor or tumor-adjacent tissues were acquired in parallel from three different instruments (Figure 1). Each omics dataset was analyzed and annotated separately. Dysregulated proteins, phosphopeptides, metabolites, and lipids were selected and summarized for both early stage and advanced stage HCC, respectively.

**Figure 1.**
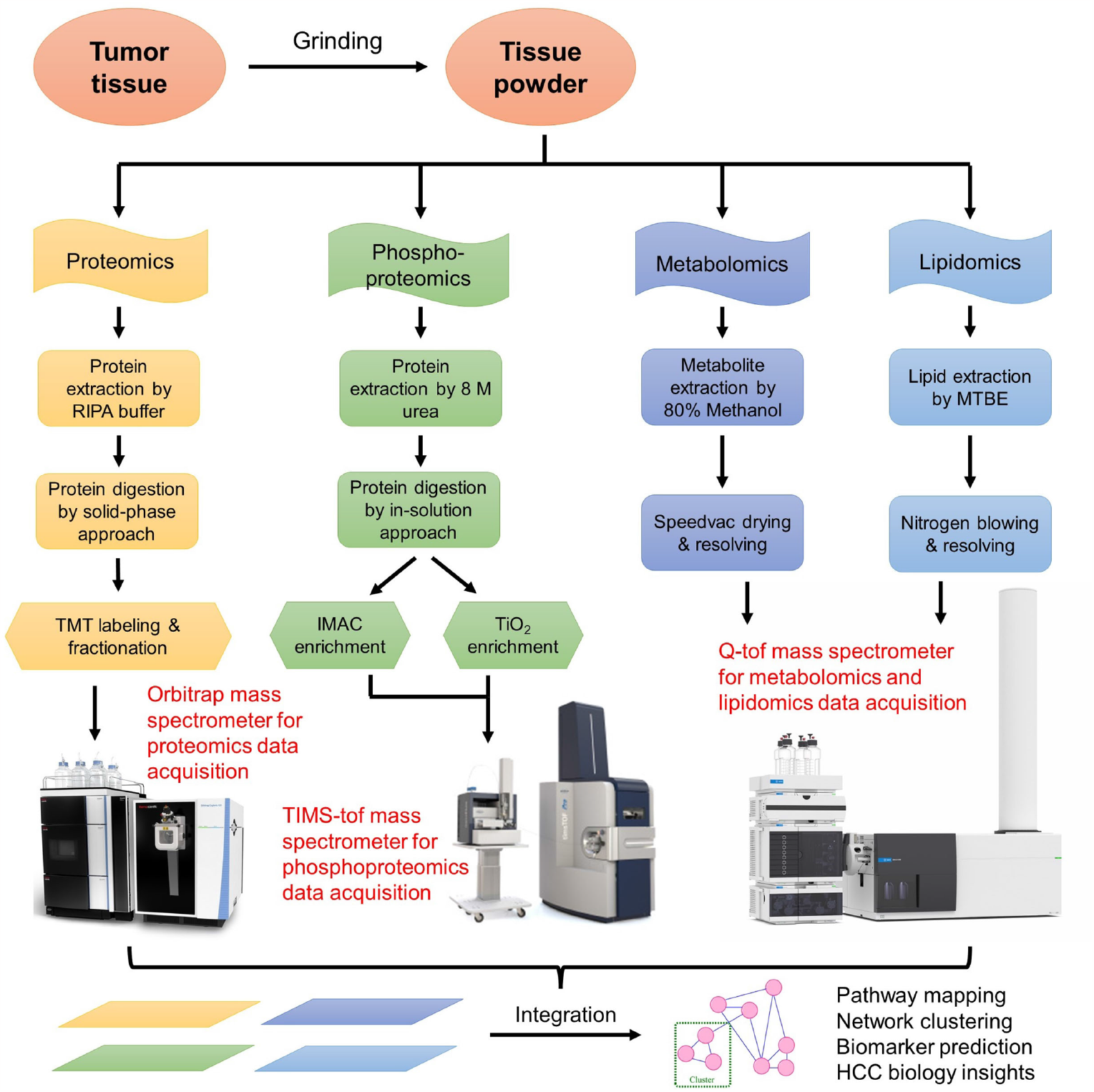
Schematic representation of multi-omic experiments including proteomics, phosphoproteomics, metabolomics and lipidomics.

### 3.1. Overview of Proteomic, Metabolomic, and Lipidomic Results

For the TMT-labeled proteomic study, a total of 12,152 protein groups (Table S1) and 166,260 unique peptides were identified from 742,816 PSMs. Among these proteins, 233 upregulated and 303 downregulated proteins were detected in early-stage HCC tissues compared to cognate tumor-adjacent tissues (Figure 2A). In contrast, 299 upregulated and 632 downregulated proteins were identified in advanced-stage HCC (Figure 2B). As expected, more dysregulated proteins were present during disease progression. GO analysis was used to cluster dysregulated proteins involved in metabolic processes, cell cycle and organization, nucleic acid and protein metabolism, transport, signal transduction, and development in both early and advanced stages (Figure S1). Analyzed by KEGG, cell cycle, DNA replication, and repair pathways were enriched in upregulated proteins, while various metabolism pathways were enriched in downregulated proteins (Figure S2).

**Figure 2.**
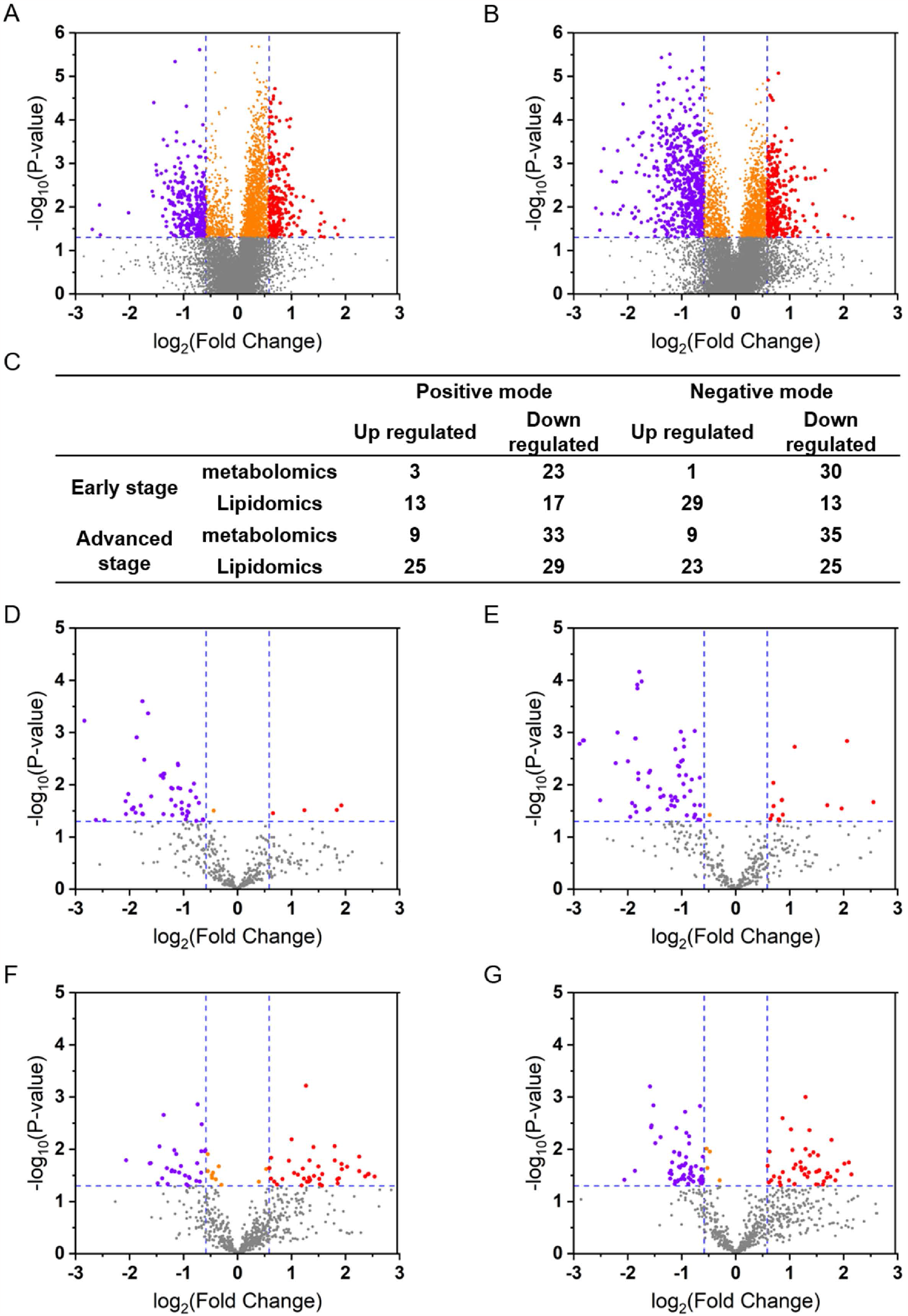
Proteomics, metabolomics and lipidomics of early and advanced-stage HCC. (A) and (B) Volcano plot analysis of TMT-labeled proteomics on early (A) and advanced (B) stage HCC. (C) The numbers of differentially accumulated metabolites and lipids quantified between early or advanced stage tumor and the normal tissue adjacent to the tumor. (D) and (E) Volcano plot analysis of metabolites quantified in early (D) or advanced (E) stage HCC. (F) and (G) Volcano plot analysis of lipids quantified in early (F) or advanced (G) stage HCC. The Statistically significantly (p-value<0.05, Student’s t-test) up-(Fold change>1.5) and down-regulated (Fold change<0.67) molecules were showed in purple and red dots, respectively.

For metabolomics and lipidomics, 251 metabolites and 330 lipids were annotated from 1,201 metabolite and 3,354 lipid features under positive mode; simultaneously, 245 metabolites and 395 lipids were identified from 1,197 metabolite and 2,698 lipid features under negative mode (Table S2). The numbers of upregulated and downregulated metabolites and lipids under positive/negative mode for both early-stage and advanced-stage HCC were summarized in Figure 2C. Combining the compounds detected from positive and negative modes, a total of 58 metabolites and 72 lipids were dysregulated in early-stage HCC, and 86 metabolites and 102 lipids were dysregulated in advanced-stage HCC (Figure 2D-2G). Consistent with the trend observed in proteomic data, the higher the tumor stage, the greater the number of significantly changed compounds discovered.

### 3.2. Phosphoproteomics and Kinase-substrates Relationship

For label-free quantitative phosphoproteomics, 25,071 unique phosphopeptides containing 35,574 unique phosphorylation sites were profiled on 6,686 proteins (Table S3). As illustrated in Figure 3A, 49.4% of phosphopeptides were identified simultaneously under both conditions, while 45.3% and 5.3% were profiled exclusively by IMAC-Fc and TiO_2_, respectively. The distribution of single-, double-, and multiple-phosphorylation sites on each phosphopeptide is shown in Figure 3B, with nearly 90% of peptides being mono-phosphorylated. The ratios of phosphorylation sites on Ser, Thr, and Tyr were exhibited in Figure 3C; consistent with basic biochemical knowledge, the ratio of detected phosphoTyosine was less than 3%. Only peptides with a single phosphorylation site were used for further quantitative variance analysis. The numbers of up- and down-regulated phosphopeptides with unique mono-phosphorylation sites in early and advanced-stage HCC were calculated and displayed on the volcano plots (Table 2 & Figure 3D-3E).

**Table 2.** The numbers of dysregulated phosphopeptides.

| Table 2. The numbers of dysregulated phosphopeptides |  |  |
| --- | --- | --- |
|  | Up-regulation | Down-regulation |
| Early-stage | 1145 | 825 |
| Advanced-stage | 1253 | 945 |

**Figure 3.**
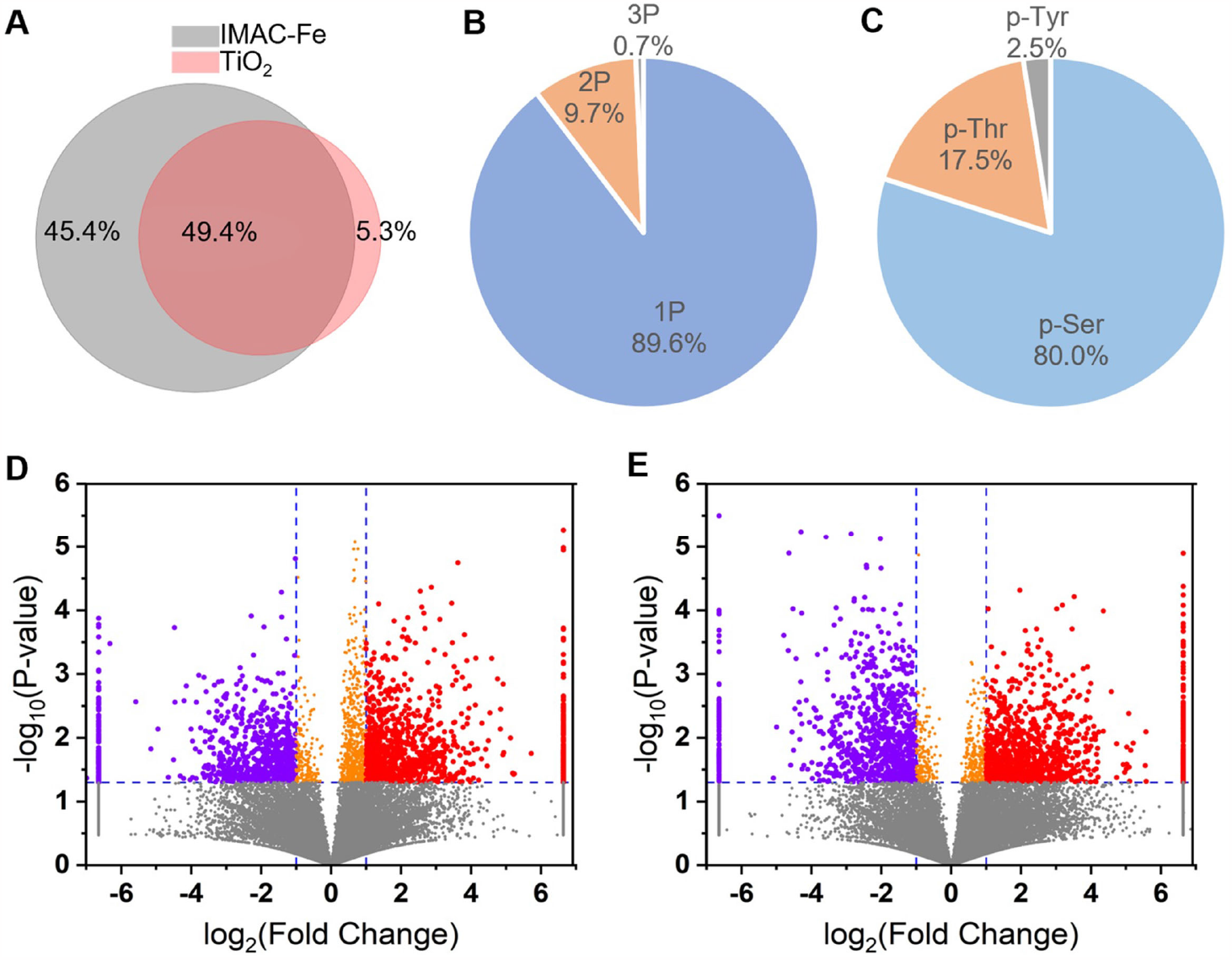
Phosphopeptide enrichment of early and advanced-stage HCC using the IMAC Fe-NTA and TiO_2_ strategies. (A) Overlap between all phosphopeptides quantified by IMAC Fe-NTA or TiO_2_ phosphopeptides enrichment method. (B) The composition of the mono-(1P), double-(2P), and multi-(3P) phosphorylated peptides quantified. (C) The percentage of phosphorylated serine (p-ser), threonine (p-thr), and tyrosine (p-tyr) in the quantified phosphopeptides. (D) Volcano plot analysis of quantified mono-phosphorylated peptides between early-stage tumor and the normal tissue adjacent to the tumor. (E) Volcano plot analysis of quantified mono-phosphorylated peptides between advanced-stage tumor and the normal tissue adjacent to the tumor. Statistically significantly (p-value<0.05, Student’s t-test) up-(Fold change>2) and down-regulated (Fold change<0.5) phosphopeptides with unique mono-phosphorylation sites were showed in purple and red dots, respectively.

The kinase-substrate relationship assay was performed using the KSEA App to predict altered kinase activities in early and advanced-stage HCC from phosphoproteomic data. The enriched kinase activities in early and advanced-stage HCC are displayed in Figure 4A and 4B, respectively. Kinases represented by red bars (z-score ≥ 2) indicate hyperactive kinase activity, while kinases denoted by blue bars (z-score ≤ -2) represent deactivated kinase activity. On one hand, most hyperactive kinase families were shared in both early and advanced-stage HCC. The high activation of cyclin-dependent kinase (CDK), mitogen-activated protein kinase (MAPK), and Aurora kinase (AURK) was expected, as they are all related to the cell cycle, cell proliferation, and mitosis, leading to tumorigenesis. On the other hand, apart from AMP-activated protein kinase (AMPK), deactivated kinases exhibited some degree of diversity, indicating distinct signal transduction pathways in different stages of HCC, which will be discussed later.

**Figure 4.**
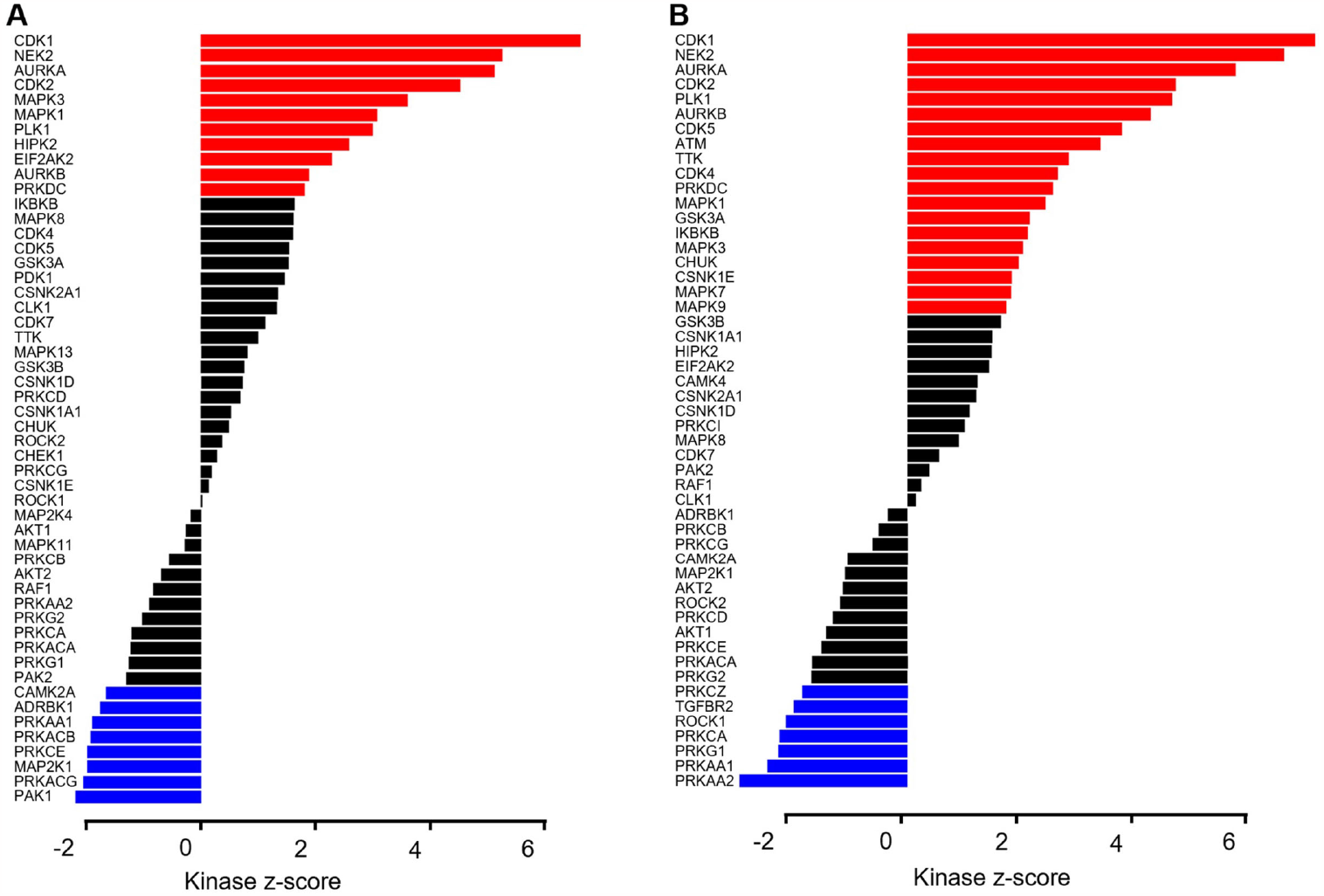
Kinase-substrate enrichment analysis of phosphoproteomic data of early-or advanced-stage HCC. (A) and (B) The enriched kinase identified in early (A) or advanced (B) stage tumor tissues when compared with the normal tissue adjacent to the tumor. Significantly hyperactivated and deactivated kinase are illustrated by red or blue, respectively. Significance thresholds were set to a z-score cutoff of 2, p value cutoff of 0.05, and substrate count cutoff of

### 3.3. Multi-Omic Characterization of Early and Advanced-Staged HCC

The lists of dysregulated proteins, metabolites, and lipids were combined and subjected to IPA for core analysis, abbreviated as PML analysis. Compared to advanced-stage HCC, a limited number of pathways were enriched in early-stage HCC, indicating that a few universal tumorigenesis processes and pathways were activated or inhibited, including DNA replication, cell proliferation, transcription factor activities, and p53 function (Figure S3). As HCC progressed to advanced stage, an increasing number of bioprocesses and pathways contributing to tumor growth were activated (Figure S4A), and various metabolism pathways, particularly those involved in anabolism, were inhibited (Figure S4B), consistent with findings from separate proteomic and metabolomic data. Two selected networks of the multi-omic assay in advanced-stage HCC are exhibited in Figure 5: one depicts a network from glucose depletion to triggering DNA damage response, while the other mostly describes how retinoid X receptor influences lipid metabolism. Although these networks were reconstructed based on the current database containing diverse information from different cell lines, tissues, and species, these findings could help researchers understand connections between individual changes in proteins or compounds and reveal the complexity of the biological system.

**Figure 5.**
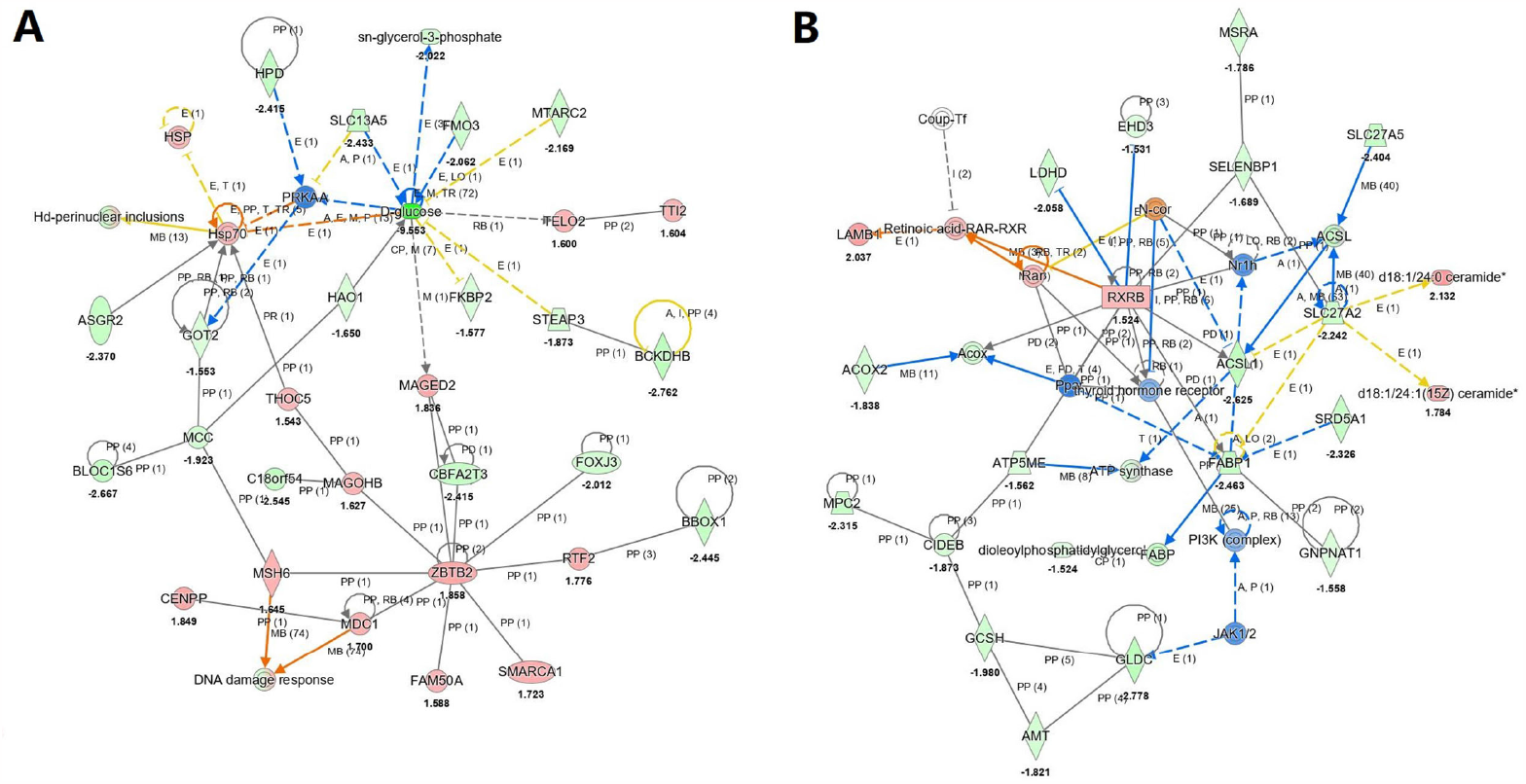
Two examples (A & B) of the protein compound interaction network based on IPA PML analysis.

Phosphorylation assay was submitted to IPA as a separate analysis. In contrast to the results of PML analysis, several kinases signaling pathways were prone to activation for tumor initiation and metastasis in early-stage HCC (Figure S5A), suggesting that post-translational modification (PTM) signaling is more sensitive than protein and metabolite changes. As expected, the bioprocesses and pathways enriched in advanced-stage HCC demonstrated a higher degree of malignancy (Figure S5B). Figure S6 displays a complex phosphorylation interaction network predicted from the phosphoproteomic data, revealing multiple protein phosphorylation factors affecting ATPase. Although phosphorylation data could not be merged with other omics data in IPA, the comparison assay provided an alternative way to integrate observations from the four omics under the current study.

### 3.4. Comparison of Advanced and Early-Stage HCC

To identify the most clinically relevant targets from dysregulated multi-omics data, we conducted a comparison of up- and down-regulated proteins and phosphoproteins using the CPTAC database [22, 29]. Figure 6A and 6B illustrate that approximately 50% of the up-regulated proteins and phosphoproteins from the CPTAC database were detected in either early or advanced-stage samples. Notably, 8 up-regulated proteins appeared in all three groups (Figure 6A), designating them as the most prominent biomarker panel. Higher expression of this gene signature was significantly associated with shorter survival (Figure 6C). Furthermore, 22 up-regulated proteins in Figure 6A were identified as specific early-stage HCC biomarkers, while 19 up-regulated proteins were specific to advanced-stage HCC (Figure 6A). Intriguingly, both gene signature groups demonstrated higher expression correlated with shorter survival, but the 19-gene signature group exhibited a longer time distance between high and low gene expression survival curves compared to the 22-gene signature group (Figure 6D & 6E). High expression of the advanced group panel indicated a longer time distance (Figure 6F), representing lower survival time compared to the high expression of the early group panel, highlighting the clinical relevance of our specific biomarker panels. A similar trend was also observed for down-regulated proteins (Figure S7).

**Figure 6.**
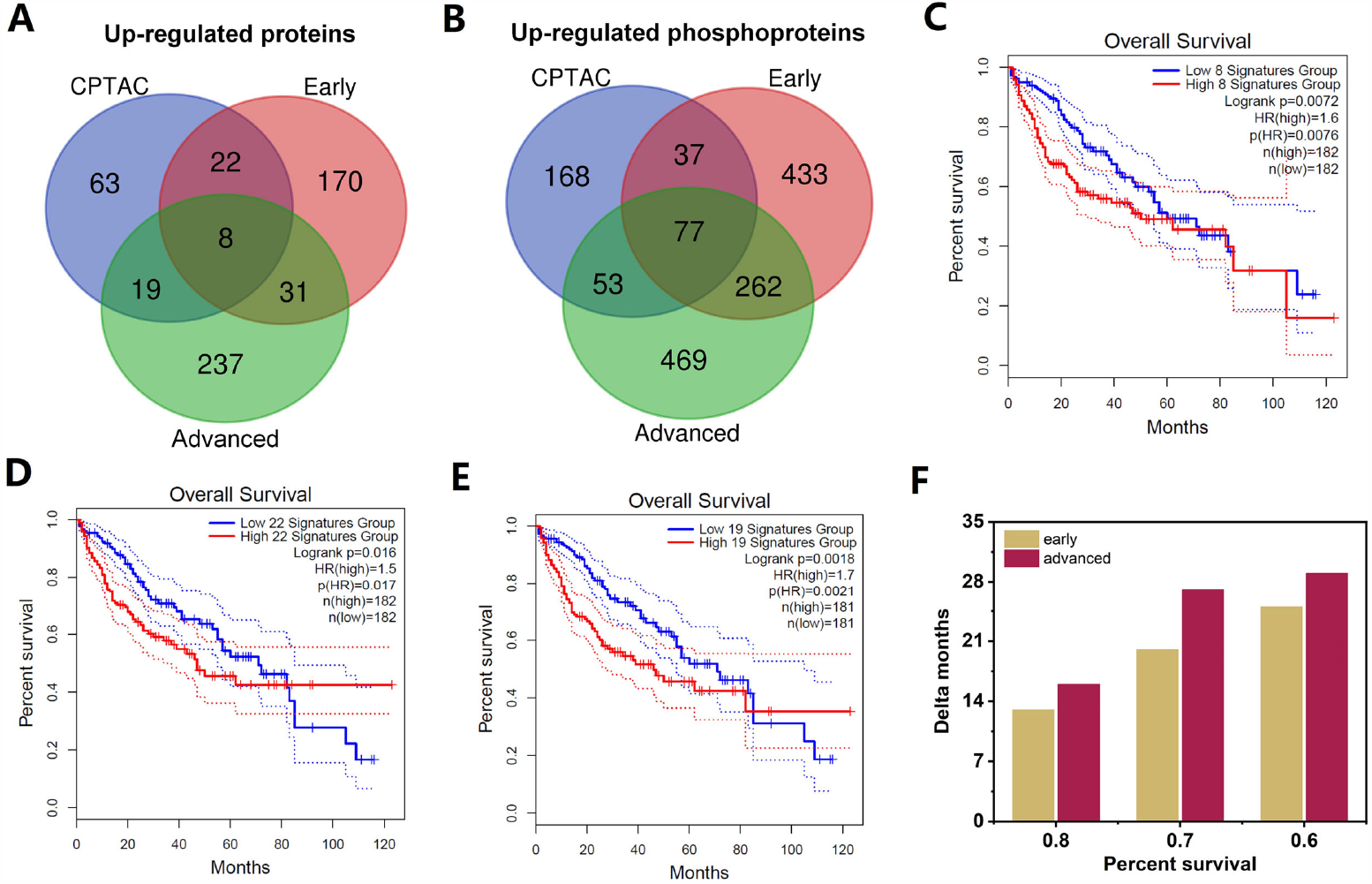
Potential biomarker panels with clinical prognostic significance. Venn diagram showing the overlap among the up-regulated proteins (A) and phosphoproteins (B) analyzed from CPTAC database, early- and advanced stage HCC data in current study. Kaplan–Meier analysis for the overall survival of the 8-(C), 22-(D) and 19-(E) gene signatures. The comparisons of the reduced month time of the high expression between early- and advanced-stage specific gene signatures (F).

By utilizing the comparison assay, the enriched a hundred pathways either activated or inhibited from both phosphorylation analysis and PML analysis were summarized in a single graphical representation (Figure 7). Most of the pathway changing trends were consistent with each other, and more significant activation or inhibition (darker colors) was observed in advanced-stages and CPTAC data than in early-stages. Some of the top changing pathways were obviously to understand. For instance, the top two enriched pathways were related to the dysregulation of xenobiotic metabolism, leading to the loss of detoxification function of liver [41]. Serotonin degradation (the third line in Figure 7), have been widely reported to be strongly inhibited, leading to serotonin accumulation, and promoting tumor growth in HCC [42]. As mentioned previously, the PML analysis in early stages only revealed some top changing pathways. In contrast, the phosphorylation analysis in early stages exposed most of the changing pathways observed in advanced stages. This phenomenon suggests that protein phosphorylation is an upstream signal of protein and small compound changes. This comparison assay list provides an overall perspective of biological pathways in HCC initiation and progression.

**Figure 7.**
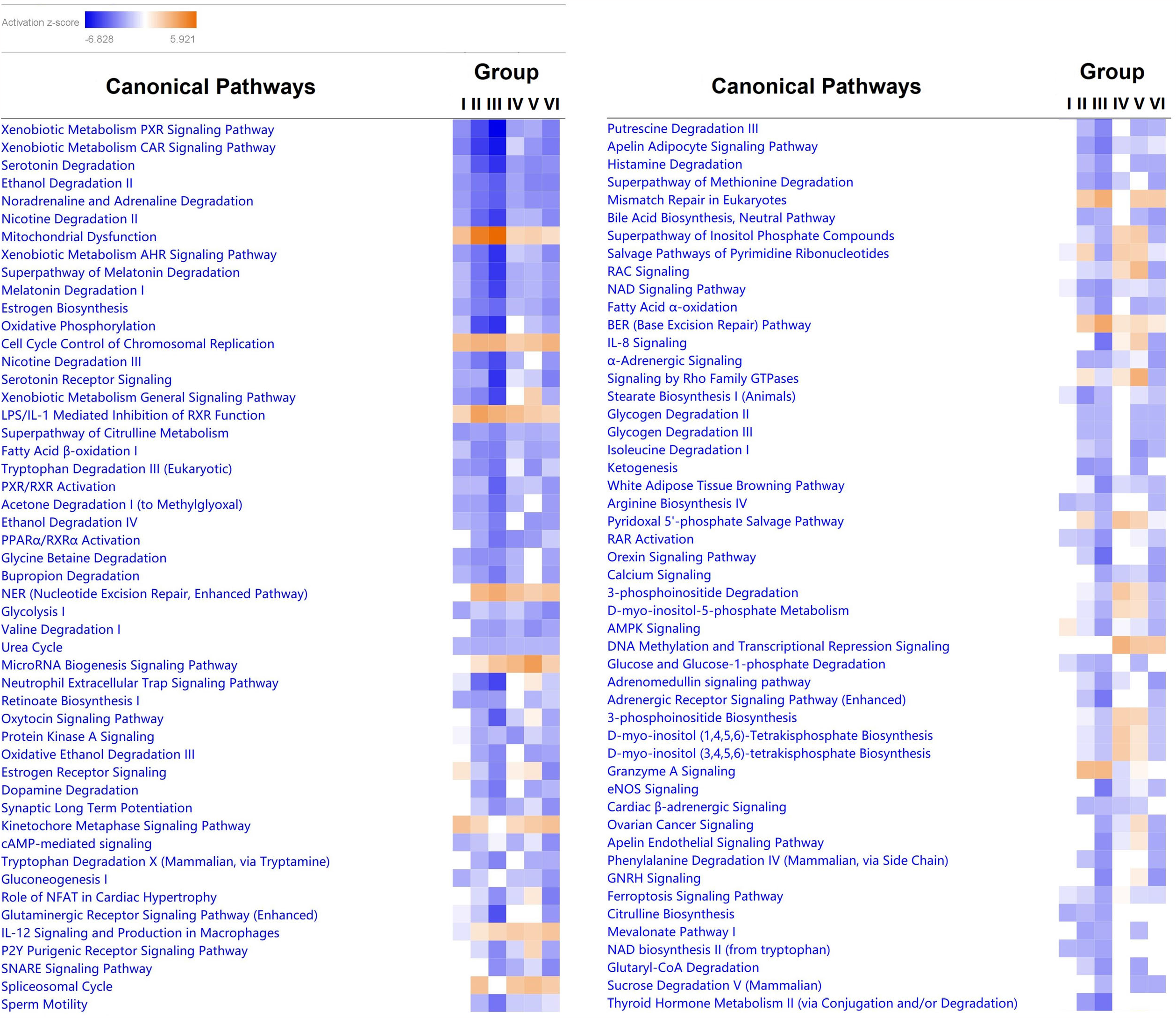
The comparison analysis of enriched pathways for PML assay in early-stage HCC (Group I), PML assay in advanced-stage HCC (Group II), proteome assay of CPTAC database (Group III), phosphorylation assay in early-stage HCC (Group IV), phosphorylation assay in advanced-stage HCC (Group V) and phosphorylation assay of CPTAC database (Group VI). The orange color and blue color in the heatmap represent pathway activation and inhibition, respectively. The darker color is related to larger z-score, which means the higher degree of activation or inhibition.

## 4. Discussion

Hepatocellular carcinoma (HCC) is one of the most malignant tumors worldwide, posing a severe threat to human health. Although surgical resection and liver transplantation can efficiently treat early-stage HCC, early diagnosing and intervening in advanced-stage HCC patients remain challenging. Multi-omic technologies offer systematic insights into the complex biological processes related to disease initiation and development, which are helpful for developing biomarkers for diagnosis and drug targets for systemic therapy. In this study, total proteins, enriched phosphopeptides, metabolites, and lipids were extracted separately from the same HCC tumor or tumor-adjacent tissues, with four omics performed on three different LC-MS/MS systems (Figure 1). Variance analysis was used to select dysregulated proteins, phosphorylated peptides, metabolites, and lipids, which were further integrated and analyzed by IPA. Distinct patterns for early-stage HCC and advanced-stage HCC were described separately (Figures 2-5), with the progression of HCC characterized by multi-omic interpretation utilizing CPTAC database (Figure 6-7). Several points related to this study warrant discussion.

Firstly, at the technical level, multi-omics and their applications are still challenging in today’s research. Simultaneously acquiring multi-omic data from the same tissue requires an adequate sample amount and diverse instrument support, especially for PTM-omics, which typically need substantial proteins and sophisticated experimental workflows, such as enrichment and fractionation. As shown in Figure 6, protein phosphorylation exhibited higher sensitivity for signal alteration than downstream proteins and compounds in early-stage HCC, suggesting that it may be more suitable for biomarker selection in early diagnosis theoretically [43-44]. Although phosphopeptides, ordinary tryptic peptides, metabolites, and lipids are all detected by LC-MS/MS, at a practical level, phosphoproteomics is considerably more complex and challenging to implement for clinical samples than other omics. Previous studies have identified over 20,000 phosphopeptides or phosphorylation sites in HCC tumors using high-pH reverse phase fractionation (one sample, six injections) and TiO_2_ enrichment [23-34]. Taking advantage of the high sensitivity and speed of the timsTOF Pro 2 mass spectrometer and the complementary results of IMAC Fe-NTA and TiO2 profiling [45-47], a comparable number of phosphopeptides has been obtained using a much simpler approach (one sample, two injections; routine DDA data acquisition without project-specific proteomics libraries). Additionally, multi-omic data have traditionally been difficult to integrate due to their complexity and heterogeneity [48]. With the assistance of the IPA platform, all the omic data generated from proteins, including PTMs, to small compounds in the current study have been recognized and integrated for analysis, revealing enriched pathways and biomarkers in different stages of HCC. All reagents, instruments, search and analysis engines used in this research are commercial or open source, reflecting the lowered threshold for multi-omic studies.

Next, the down-regulated proteins and metabolites from proteomics and metabolomics were found to be quite consistent, demonstrating a global attenuation of cell metabolism that prominently occurred in advanced-stage HCC. Unlike the phenotypes obtained from cell lines or mouse models, both the enzymes and metabolites in glycolysis and anaerobic respiration in these clinical samples showed significant decreases, seemingly opposite to the Warburg effect. However, MAPK deactivation from phosphoproteomic data definitively supported the activation of aerobic glycolysis [49-50]. The paradox can be explained from several perspectives. As mentioned earlier, proteome and metabolome changes are the downstream consequence of cell signaling, and the tumor tissue may already be in a low-glucose equilibrium if insufficient materials are supplied. Additionally, the traditional Warburg effect has been increasingly challenged due to different cancer types and diverse tumor microenvironments, and these phenomena might also be caused by tumor stromal cells, resulting in the observation of a reverse Warburg effect [51-53]. Moreover, glutamate was significantly increased in advanced-stage HCC, indicating the activation of an alternative energy generation pathway. Although the glutamine pathway can compensate for some energy supply, it still seemed that global catabolism decreases during tumor progression. This viewpoint has been recently demonstrated by a Flux work that indicates solid tumors generally produce ATP at a slower rate than normal [54].

Finally, the multi-omic integrated assay revealed comprehensive signal networks, providing new insights into the mechanisms of HCC disease. As described above, most reported biological processes in HCC, including epithelial-mesenchymal transition (EMT), abnormal tumor microenvironment formation, and senescence bypass, emerged through the multi-omic data. EMT is considered one of the common features of cancer development [55], and in HCC, besides transcription factor promotion, signaling pathways such as AURK, MAPK, and NF-κB also play important roles in the EMT process [56-58]. Consistent with previous studies, these canonical kinase pathways were hyperactive from early to advanced-stage HCC. Our data also contained some differing results that seemed contradictory to previous reports; for example, Rho-associated coiled-coil-containing protein kinase 1 (ROCK1) was significantly deactivated in advanced-stage HCC. Although ROCK2 was observed as an overexpressed kinase leading to tumor malignancy both by proteomic profiling and biology studies [23, 59], ROCK1 might contribute to HCC progression through an alternative way involving the peritumoral microenvironment (PME), a new concept describing the tissue surrounding the tumor that influences cancer development [60]. Existing evidence showed that hepatic stellate cells (HSCs) accumulating in HCC PME with an increase in VEGF were able to induce angiogenesis [61], and ROCK1 was reported as a negative regulator of VEGF-driven angiogenic activation [62]. We hypothesize that deactivated ROCK1 would appear in HSCs and further promote angiogenesis. Due to the heterogeneity of tumor tissue, omic studies in this research could not identify specific cell types. Although this limitation might be partially addressed by bioinformatic assays [25], elaborate biological experiments are needed for validation. Additionally, glycogen synthase kinase 3α (GSK3A) was hyperactivated in advanced-stage HCC, which is responsible for glycogen accumulation and tumorigenesis by the suppression of Hippo signaling through glycogen-induced phase separation [63]. The simultaneous discovery of ROCK1 deactivation and GSK3A activation only in advanced-stage HCC adds new insights to the HCC progression landscape.

In conclusion, performing multi-omic profiling from proteins to small compounds provided vast amounts of information in early and advanced-stage HCC. We demonstrated a feasible and comprehensive workflow for multi-omic integration studies, encompassing experimental design, sample preparation, data acquisition, and analysis. We also established a correlation network of HCC initiation and development, proposing new mechanisms for HCC, including reversed hypermetabolism and angiogenesis promoted by ROCK1 deactivation. Though the HCC sample size was quite limited in the current study, the parameters in each group, e.g. gender, alcohol drinking, etc., were fairly close [64]. Our results can also be verified by CPTAC data to some extent, these insights can broaden our knowledge pool and provide research directions to cancer biologists.

## Supporting information

supplementary part

## Author Contributions

Conceptualization, S.F. and W.X.; methodology, J.H., M.F. and W.X.; formal analysis, J.H., M.F. and X.X.; investigation, M.F., J.H., J.C., X.X., W.Z., X.Z. and J.P.; data curation, J.C.; writing-original draft preparation, S.F.; writing-review and editing, M.F, S.F. and W.Z. All authors have read and agreed to the published version of the manuscript.

## Institutional Review Board Statement

The study was conducted according to the guidelines of the Declaration of Helsinki, and approved by the Research Ethics Committee of Shulan (Hangzhou) Hospital (Reference Number: KY2023033).

## Data Availability Statement

The datasets used and/or analyzed during the current study are available at the Massive website (https://massive.ucsd.edu/ProteoSAFe/static/massive.jsp) with the ID number of MSV000091782.

## Acknowledgments

We thank Dr. Yalin Wang, the director of the Biomedical Research Core Facility at Westlake University for supporting this research. We also thank Beibei Ma and Xiaoxian Du from Bruker Corporation for the database searching and quantification of the MS data acquired by the timsTOF Pro2 mass spectrometer.

## References

1. Forner, A.; Llovet, J.M.; Bruix, J. Hepatocellular carcinoma. Lancet. 2012, 379, 1245–1255.

2. Hartke, J.; Johnson, M.; Ghabril, M. The diagnosis and treatment of hepatocellular carcinoma. Semin Diagn Pathol. 2017, 34, 153–159.

3. Llovet, J.M.; Kelley, R.K.; Villanueva, A.; Singal, A.G.; Pikarsky, E.; Roayaie, S.; Lencioni, R.; Koike, K.; Zucman-Rossi, J.; Finn R.S. Hepatocellular carcinoma. Nat Rev Dis Primers. 2021, 7, 6.

4. Ganesan, P.; Kulik, L.M. Hepatocellular Carcinoma: New Developments. Clin Liver Dis. 2023, 27, 85–102.

5. Sung, H.; Ferlay, J.; Siegel, R.L.; Laversanne, M.; Soerjomataram, I.; Jemal, A.; Bray, F. Global Cancer Statistics 2020: GLOBOCAN Estimates of Incidence and Mortality Worldwide for 36 Cancers in 185 Countries. CA Cancer J Clin. 2021, 71, 209–249.

6. Ding, J.; Wen, Z. Survival improvement and prognosis for hepatocellular carcinoma: analysis of the SEER database. BMC Cancer. 2021, 21, 1157.

7. Piñero, F.; Dirchwolf, M.; Pessôa, M.G. Biomarkers in Hepatocellular Carcinoma Diagnosis, Prognosis and Treatment Response Assessment. Cells. 2020, 9, 1370.

8. Galle, P.R.; Foerster, F.; Kudo, M.; Chan, S.L.; Llovet, J.; Qin, S.; Schelman, W.; Chintharlapalli, S.; Abada, P.; Sherman, M.; et al. Biology and significance of alpha-fetoprotein in hepatocellular carcinoma. Liver Int. 2019, 39, 2214–2229.

9. Grandhi, M.S.; Kim, A.K.; Ronnekleiv-Kelly, S.M.; Kamel, I.R.; Ghasebeh, M.A.; Pawlik, T.M. Hepatocellular carcinoma: From diagnosis to treatment. Surg Oncol. 2016, 25, 74–85.

10. Tellapuri, S.; Sutphin, P.D.; Beg, M.S.; Singal, A.G.; Kalva, S.P. Staging systems of hepatocellular carcinoma: A review. Indian J Gastroenterol. 2018, 37, 481–491.

11. Marrero, J.A.; Kulik, L.M.; Sirlin, C.B.; Zhu, A.X.; Finn, R.S.; Abecassis, M.M.; Roberts, L.R.; Heimbach, J.K. Diagnosis, Staging, and Management of Hepatocellular Carcinoma: 2018 Practice Guidance by the American Association for the Study of Liver Diseases. Hepatology. 2018, 68, 723–750.

12. Fan, J.; Yang, G.S.; Fu, Z.R.; Peng, Z.H.; Xia, Q.; Peng, C.H.; Qian, J.M.; Zhou, J.; Xu, Y.; Qiu, S.J.; et al. Liver transplantation outcomes in 1,078 hepatocellular carcinoma patients: a multi-center experience in Shanghai, China. J Cancer Res Clin Oncol. 2009, 135, 1403–1412.

13. Tateishi, R.; Shiina, S.; Teratani, T.; Obi, S.; Sato, S.; Koike, Y.; Fujishima, T.; Yoshida, H.; Kawabe, T.; Omata, M. Percutaneous radiofrequency ablation for hepatocellular carcinoma. An analysis of 1000 cases. Cancer. 2005, 103, 1201–1209.

14. Llovet, J.M.; Castet, F.; Heikenwalder, M.; Maini, M.K.; Mazzaferro, V.; Pinato, D.J.; Pikarsky, E.; Zhu, A.X.; Finn, R.S. Immunotherapies for hepatocellular carcinoma. Nat Rev Clin Oncol. 2022, 19, 151–172.

15. Llovet, J.M.; Ricci, S.; Mazzaferro, V.; Hilgard, P.; Gane, E.; Blanc, J.F.; de Oliveira, A.C.; Santoro, A.; Raoul, J.L.; Forner, A.; et al. Sorafenib in advanced hepatocellular carcinoma. N Engl J Med. 2008, 24, 378–390.

16. Keating, M. G.; Santoro, A. Sorafenib: a review of its use in advanced hepatocellular carcinoma. Drugs. 2009, 69, 223–240.

17. Yarchoan, M.; Agarwal, P.; Villanueva, A.; Rao, S.; Dawson, L.A.; Llovet, J.M.; Finn, R.S.; Groopman, J.D.; El-Serag, H.B.; Monga, S.P.; et al. Recent Developments and Therapeutic Strategies against Hepatocellular Carcinoma. Cancer Res. 2019, 79, 4326–4330.

18. Finn, R.S.; Qin, S.; Ikeda, M.; Galle, P.R.; Ducreux, M.; Kim, T.Y.; Kudo, M.; Breder, V.; Merle, P.; Kaseb, A.O. et al. Atezolizumab plus Bevacizumab in Unresectable Hepatocellular Carcinoma. N Engl J Med. 2020, 382, 1894–1905.

19. Schulze, K.; Nault, J.C.; Villanueva, A. Genetic profiling of hepatocellular carcinoma using next-generation sequencing. J Hepatol. 2016, 65, 1031–1042.

20. Jin, Y.; Lee, W.Y.; Toh, S.T.; Tennakoon, C.; Toh, H.C.; Chow, P.K.; Chung, A.Y.; Chong, S.S.; Ooi, L.L.; Sung, W.K.; et al. Comprehensive analysis of transcriptome profiles in hepatocellular carcinoma. J Transl Med. 2019, 17, 273.

21. Megger, D.A.; Naboulsi, W.; Meyer, H.E.; Sitek, B. Proteome Analyses of Hepatocellular Carcinoma. J Clin Transl Hepatol. 2014, 2, 23–30.

22. Wu, Z.H.; Yang, D.L. Identification of a protein signature for predicting overall survival of hepatocellular carcinoma: a study based on data mining. BMC Cancer. 2020, 20, 720.

23. Ren, L.; Li, C.; Wang, Y.; Teng, Y.; Sun, H.; Xing, B.; Yang, X.; Jiang, Y.; He, F. In Vivo Phosphoproteome Analysis Reveals Kinome Reprogramming in Hepatocellular Carcinoma. Mol Cell Proteomics. 2018, 17, 1067–1083.

24. Jiang, Y.; Sun, A.; Zhao, Y.; Ying, W.; Sun, H.; Yang, X.; Xing, B.; Sun, W.; Ren, L.; Hu, B.; et al. Proteomics identifies new therapeutic targets of early-stage hepatocellular carcinoma. Nature. 2019, 567, 257–261.

25. Gu, Y.; Guo, Y.; Gao, N.; Fang, Y.; Xu, C.; Hu, G.; Guo, M.; Ma, Y.; Zhang, Y.; Zhou, J.; et al. The proteomic characterization of the peritumor microenvironment in human hepatocellular carcinoma. Oncogene. 2022, 41, 2480–2491.

26. Wu, X.; Xing, X.; Dowlut, D.; Zeng, Y.; Liu, J.; Liu, X. Integrating phosphoproteomics into kinase-targeted cancer therapies in precision medicine. J Proteomics. 2019, 191, 68–79.

27. Moldogazieva, N.T.; Mokhosoev, I.M.; Zavadskiy, S.P.; Terentiev, A.A. Proteomic Profiling and Artificial Intelligence for Hepatocellular Carcinoma Translational Medicine. Biomedicines. 2021, 9, 159.

28. Mani, DR.; Krug, K.; Zhang, B.; Satpathy, S.; Clauser, K.R.; Ding, L.; Ellis, M.; Gillette, M.A.; Carr, S.A. Cancer proteogenomics: current impact and future prospects. Nat Rev Cancer. 2022, 22, 298–313.

29. Gao, Q.; Zhu, H.; Dong, L.; Shi, W.; Chen, R.; Song, Z.; Huang, C.; Li, J.; Dong, X.; Zhou, Y.; et al. Integrated Proteogenomic Characterization of HBV-Related Hepatocellular Carcinoma. Cell. 2019, 179, 561–577.

30. Ng, C.K.Y.; Dazert, E.; Boldanova, T.; Coto-Llerena, M.; Nuciforo, S.; Ercan, C.; Suslov, A.; Meier, M.A.; Bock, T.; Schmidt, A.; et al. Integrative proteogenomic characterization of hepatocellular carcinoma across etiologies and stages. Nat Commun. 2022, 13, 2436.

31. Khalil, A.; Elfert, A.; Ghanem, S.; Helal, M.; Abdelsattar, S.; Elgedawy, G.; Obada, M.; Abdel-Samiee, M.; El-Said, H. The role of metabolomics in hepatocellular carcinoma. Egyptian Liver J. 2021, 11, 41.

32. Feng, N.; Yu, F.; Yu, F.; Feng, Y.; Zhu, X.; Xie, Z.; Zhai, Y. Metabolomic biomarkers for hepatocellular carcinoma: A systematic review. Medicine (Baltimore). 2022, 101, e28510.

33. Ferrarini, A.; Di Poto, C.; He, S.; Tu, C.; Varghese, R.S.; Kara Balla, A.; Jayatilake, M.; Li, Z.; Ghaffari, K.; Fan, Z.; et al. Metabolomic Analysis of Liver Tissues for Characterization of Hepatocellular Carcinoma. J Proteome Res. 2019, 18, 3067–3076.

34. Sangineto, M.; Villani, R.; Cavallone, F.; Romano, A.; Loizzi, D.; Serviddio, G. Lipid Metabolism in Development and Progression of Hepatocellular Carcinoma. Cancers (Basel). 2020, 12, 1419.

35. Tan, S.L.W.; Israeli, E.; Ericksen, R.E.; Chow, P.K.H.; Han, W. The altered lipidome of hepatocellular carcinoma. Semin Cancer Biol. 2022, 86, 445–456.

36. Beyoğlu, D.; Idle, J.R. Metabolomic and Lipidomic Biomarkers for Premalignant Liver Disease Diagnosis and Therapy. Metabolites. 2020, 10, 50.

37. Ashburner, M.; Ball, C.A.; Blake, J.A.; Botstein, D.; Butler, H.; Cherry, J.M.; Davis, A.P.; Dolinski, K.; Dwight, S.S.; Eppig, JT.; et al. Gene ontology: tool for the unification of biology. The Gene Ontology Consortium. Nat Genet. 2000, 25, 25–29.

38. Mi, H.; Muruganujan, A.; Ebert, D.; Huang, X.; Thomas, P.D. PANTHER version 14: more genomes, a new PANTHER GO-slim and improvements in enrichment analysis tools. Nucleic Acids Res. 2019, 47, D419–D426.

39. Moriya, Y.; Itoh, M.; Okuda, S.; Yoshizawa, A.C.; Kanehisa, M. KAAS: an automatic genome annotation and pathway reconstruction server. Nucleic Acids Res. 2007, 35, W182–185.

40. Wiredja, D. D.; Koyutürk, M.; Chance, M. R. The KSEA App: a web-based tool for kinase activity inference from quantitative phosphoproteomics. Bioinformatics (Oxford, England), 2017, 33, 3489–3491.

41. Timsit, Y.E.; Negishi, M. CAR and PXR: the xenobiotic-sensing receptors. Steroids. 2007, 72. 231–246.

42. Soll, C.; Jang, J.H.; Riener, M.O.; Moritz, W.; Wild, P.J.; Graf, R.; Clavien, P.A. Serotonin promotes tumor growth in human hepatocellular cancer. Hepatology. 2010, 51, 1244–1254.

43. Carter, A.M.; Tan, C.; Pozo, K.; Telange, R.; Molinaro, R.; Guo, A.; De Rosa, E.; Martinez, J.O.; Zhang, S.; Kumar, N.; et al. Phosphoprotein-based biomarkers as predictors for cancer therapy. Proc Nal Acad Sci. 2020, 117, 18401–18411.

44. Henriques, A.G.; Müller, T.; Oliveira, J.M.; Cova, M.; da Cruz E Silva, C.B.; da Cruz E Silva, O.A. Altered protein phosphorylation as a resource for potential AD biomarkers. Sci Rep. 2016, 6, 30319.

45. Meier, F.; Brunner, A.D.; Koch, S.; Koch, H.; Lubeck, M.; Krause, M.; Goedecke, N.; Decker, J.; Kosinski, T.; Park, M.A.; et al. Online Parallel Accumulation-Serial Fragmentation (PASEF) with a Novel Trapped Ion Mobility Mass Spectrometer. Mol Cell Proteomics. 2018, 17, 2534–2545.

46. Oliinyk, D.; Meier, F. Ion mobility-resolved phosphoproteomics with dia-PASEF and short gradients. Proteomics. 2023, 23, e2200032.

47. Yue, X.; Schunter, A.; Hummon, A.B. Comparing multistep immobilized metal affinity chromatography and multistep TiO2 methods for phosphopeptide enrichment. Anal Chem. 2015, 87, 8837–8844.

48. Subramanian, I.; Verma, S.; Kumar, S.; Jere, A.; Anamika, K. Multi-omics Data Integration, Interpretation, and Its Application. Bioinform Biol Insights. 2020, 14, 1177932219899051.

49. Steinberg, G.R.; Carling, D. AMP-activated protein kinase: the current landscape for drug development. Nat Rev Drug Discov. 2019, 18, 527–551.

50. Meng, S.S.; Gu, H.W.; Zhang, T.; Li, Y.S.; Tang, H.B. Gradual deterioration of fatty liver disease to liver cancer via inhibition of AMPK signaling pathways involved in energy-dependent disorders, cellular aging, and chronic inflammation. Front Oncol. 2023, 13, 1099624.

51. Lee, M.; Yoon, J.H. Metabolic interplay between glycolysis and mitochondrial oxidation: The reverse Warburg effect and its therapeutic implication. World J Biol Chem. 2015, 6, 148–161.

52. Chen, H.; Wu, Q.; Peng, L.; Cao, T.; Deng, M.; Liu, Y.; Huang, J.; Hu, Y.; Fu, N.; Zhou, K.; et al. Mechanism, Clinical Significance, and Treatment Strategy of Warburg Effect in Hepatocellular Carcinoma. J Nanomaterials. 2021, 2021, 5164100.

53. Feng, J.; Li, J.; Wu, L.; Yu, Q.; Ji, J.; Wu, J.; Dai, W.; Guo, C. Emerging roles and the regulation of aerobic glycolysis in hepatocellular carcinoma. J Exp Clin Cancer Res. 2020, 39, 126.

54. Bartman, C.R.; Weilandt, D.R.; Shen, Y.; Lee, W.D.; Han, Y.; TeSlaa, T.; Jankowski, C.S.R.; Samarah, L.; Park, N.R.; da Silva-Diz, V.; et al. Slow TCA flux and ATP production in primary solid tumours but not metastases. Nature. 2023, 614, 349–357.

55. Ye, X.; Weinberg, R.A. Epithelial-Mesenchymal Plasticity: A Central Regulator of Cancer Progression. Trends Cell Biol. 2015, 25, 675–686.

56. Du, R.; Huang, C.; Liu, K.; Li, X.; Dong, Z. Targeting AURKA in Cancer: molecular mechanisms and opportunities for Cancer therapy. Mol Cancer. 2021, 20, 15.

57. Chen, L.; Guo, P.; He, Y.; Chen, Z.; Chen, L.; Luo, Y.; Qi, L.; Liu, Y.; Wu, Q.; Cui, Y.; et al. HCC-derived exosomes elicit HCC progression and recurrence by epithelial-mesenchymal transition through MAPK/ERK signaling pathway. Cell Death Dis. 2018, 9, 513.

58. Shi, C.; Chen, Y.; Chen, Y.; Yang, Y.; Bing, W.; Qi, J. CD4+ CD25+ regulatory T cells promote hepatocellular carcinoma invasion via TGF-β1-induced epithelial-mesenchymal transition. Onco Targets Ther. 2018, 12, 279–289.

59. Wong, C.C.; Wong, C.M.; Tung, E.K.; Man, K.; Ng, I.O. Rho-kinase 2 is frequently overexpressed in hepatocellular carcinoma and involved in tumor invasion. Hepatology. 2009, 49, 1583–1594.

60. McGranahan, N.; Swanton, C. Clonal Heterogeneity and Tumor Evolution: Past, Present, and the Future. Cell. 2017, 168, 613–628.

61. Lu, Y.; Lin, N.; Chen, Z.; Xu, R. Hypoxia-induced secretion of platelet-derived growth factor-BB by hepatocellular carcinoma cells increases activated hepatic stellate cell proliferation, migration and expression of vascular endothelial growth factor-A. Mol Med Rep. 2015, 11, 691–697.

62. Kroll, J.; Epting, D.; Kern, K.; Dietz, C.T.; Feng, Y. Hammes, H.P.; Wieland, T.; Augustin, H.G. Inhibition of Rho-dependent kinases ROCK I/II activates VEGF-driven retinal neovascularization and sprouting angiogenesis. Am J Physiol Heart Circ Physiol. 2009, 296, H893–899.

63. Liu, Q.; Li, J.; Zhang, W.; Xiao, C.; Zhang, S.; Nian, C.; Li, J.; Su, D.; Chen, L.; Zhao, Q.; et al. Glycogen accumulation and phase separation drives liver tumor initiation. Cell. 2021, 184, 5559-5576.e19.

64. Li, H.; Rong, Z.; Wang, H.; Zhang, N.; Pu, C.; Zhao, Y.; Zheng, X.; Lei, C.; Liu, Y.; Luo, X.; et al. Wang J. Proteomic analysis revealed common, unique and systemic signatures in gender-dependent hepatocarcinogenesis. Biol Sex Differ. 2020, 11, 46.

