## supplementary part for "Comprehensive Multi-Omic Analysis Reveals Distinct Molecular Features in Early and Advanced Stages of Hepatocellular Carcinoma"

**Materials and Methods**

**1. Reagents**

Reagents used in this study included Trypsin Gold from Promega, Urea, DL-dithiothreitol (DTT), iodoacetamide (IAM), 1M triethylamonium bicarbonate (TEAB) buffer, ammonium hydroxide (30%), trifluoroacetic acid (TFA), 5M ammonium acetate solution, and ammonium fluoride from Sigma-Aldrich. Protease inhibitor cocktail for general use (100X) and Phosphatase inhibitor cocktail A (50X) were obtained from Beyotime. Pierce™ BCA Protein Assay Kit (catalog no. 23225), TMT10plex™ Label Reagent Set (catalog no. 90110), High-Select™ Fe-NTA Phosphopeptide Enrichment Kit (catalog no. A32992), Pierce TiO_2_ Phosphopeptide Enrichment and Clean-up Kit (catalog no. A32993), Hydroxylamine (50%), and formic acid (FA) were purchased from Thermo Fisher Scientific. SpeedBeads magnetic carboxylate-modified particles were sourced from Cytiva, while Methyl tert-butyl ether (MTBE) was obtained from Aladdin. All mobile phases used in this study, including water, 2-propanol, acetonitrile, methanol, and methylene chloride, were of LC-MS grade and obtained from Fisher Chemical. All other reagents were locally sourced analytical-grade products.

**2. Protein extraction, digestion, TMT labeling and fractionation**

To extract proteins, 3 mg of the ground tissue was lysed in 200 μL of RIPA buffer containing a protease inhibitor cocktail, followed by sonication for ten cycles with a 30s on and 30s off using Bioruptor Pico (Diagenode). The suspension was then centrifuged at 12,000 g for 30 min at 4ºC, and the supernatant containing proteins was transferred to a new tube. The protein concentration was determined using the BCA method [1]. Next, 50 μg of protein from each sample was digested with trypsin using the SP3 protocol [2]. Briefly, the protein concentration was adjusted to 0.5 mg/mL, followed by reduction with 10 mM DTT and alkylation with 25 mM iodoacetamide. The mixture was then incubated at room temperature at 1,000 rpm for 10 min after adding 10 μL of 50 μg/μl pre-washed beads and 100 μL of ethanol to promote binding. The solution was removed using a magnetic rack, and the beads were washed four times with 500 μL of 80% ethanol. Bound proteins were digested by adding 100 μL of 0.1 M TEAB with 0.5 μg of trypsin and incubating at 37ºC overnight. Each tryptic peptide sample was labeled with TMT10plex reagent for 1 h at room temperature (see Table S1), and the reaction was quenched by adding 8 μL of 5% hydroxylamine. TMT-labeled peptides from different channels were combined, desalted using an Oasis HLB column, and lyophilized in a speedvac.

The TMT-labeled peptides were resuspended in 100 μL of 50 mM NH_3_·H_2_O for fractionation. High-pH reverse-phase fractionation was performed using a Waters BEH C18 column (300 Å, 5 μm, 4.6×250 mm) on an Agilent 1260-Bio HPLC instrument, with the mobile phase and gradient composition detailed in reference [3]. Ten subfractions were generated from each TMT combined sample, lyophilized in a speedvac, reconstructed in 0.1% FA, and subjected to LC-MS/MS for proteomic data acquisition.

**3. Phosphoproteomics sample preparation**

Eighty mg of ground tissue was lysed in 2.5 mL of 8 M urea containing protease and phosphatase inhibitor cocktails. Protein was extracted and quantified as described previously. 5 mg protein from each sample was reduced, alkylated, and then digested in 1.5 M urea with 50 μg trypsin at 37ºC overnight. Tryptic peptide was desalted using an Oasis HLB column. 3 mg and 1.5 mg peptides were proceeded for phosphopeptide enrichment using IMAC Fe-NTA and TiO_2_ phosphopeptide enrichment kits, respectively, following the kit instructions. Phosphopeptides were eluted, desalted, lyophilized in a speedvac, and analyzed separately by LC-MS/MS.

**4. Metabolite extraction**

For metabolite extraction, pre-chilled 80% HPLC-grade methanol (500 μL) was added to 50 mg of ground tissue, followed by vertexing for 2 min on ice. The tissue suspension was then centrifuged at 12,000 g for 20 min at 4ºC, and the metabolite-containing supernatant was transferred to a new tube. Extracted metabolites were dried in a speedvac, reconstructed in 80% acetonitrile, and subjected to LC-MS/MS for metabolomic data acquisition.

**5. Lipid extraction**

For lipid extraction, 50 mg of ground tissue was resuspended in 0.2 mL of MS-grade water and 1.5 mL of HPLC-grade methanol. After vertexing for 2 min, 5 mL of MTBE was added, and the mixture was incubated at room temperature in a shaker at 300 rpm for 1 h. Phase separation was induced by adding 1.25 mL of MS-grade water, followed by centrifugation at 1,000 g for 10 min. The upper (organic) phase was collected, and the lower phase was re-extracted with 2 mL of the solvent mixture composed of MTBE:methanol:water at a ratio of 10:3:2.5. The combined organic phases were dried using a nitrogen evaporator. The extracted lipid was reconstituted in a chloroform/methanol (2:1) solution and subjected to LC-MS/MS for lipidomic data acquisition.

**6. LC-MS/MS for proteomics**

The TMT-labeled peptides were separated by a 135-min gradient elution at a flow rate of 0.300 µL/min using a Thermo Vanquish Neo integrated nano-HPLC system directly interfaced with a Thermo Exploris 480 mass spectrometer equipped with FAIMS Pro. The analytical column was a home-made fused silica capillary column (75 µm ID, 150 mm length; Upchurch, Oak Harbor, WA) packed with C-18 resin (300 Å, 2 µm, Varian, Lexington, MA). Mobile phase A consisted of 0.1% formic acid in water, and mobile phase B consisted of 80% acetonitrile and 0.1% formic acid. The mass spectrometer was operated in data-dependent acquisition mode using Xcalibur 4.1 software, and the -45 V and -65 V CV values of FAIMS Pro were set as the two independent acquisition events. Under one event, a single full-scan mass spectrum in the Orbitrap (350-1800 m/z, 60,000 resolution) was followed by several data-dependent MS/MS scans (110-1500 m/z, 15,000 resolution) at 30% normalized collision energy, with a total cycle time of 1 s per event. The AGC target was set as 5e4, and the maximum injection time was 50 ms. The Turbo TMT option was enabled. Each mass spectrum was analyzed using the Thermo Xcalibur Qual Browser and Proteome Discoverer (2.5.0.400) for the database searching against the Homo sapiens proteome database downloaded from UniProtKB (UP000005640) containing 80,581 proteins as of October 18, 2022 and TMT quantification. The Sequest search parameters included a 10 ppm precursor mass tolerance, 0.02 Da fragment ion tolerance, and up to 2 internal cleavage sites. Fixed modifications included cysteine alkylation and TMT10 modification, and the methionine oxidation was variable modification. Peptides were filtered with 1% false discovery rate (FDR).

**7. LC-MS/MS for phosphoproteomics**

The enriched phosphopeptides were separated by a 120-min gradient elution at a flow rate of 0.300 µL/min using a Bruker Nano Elute integrated nano-HPLC system directly interfaced with a Bruker timsTOF Pro 2 mass spectrometer. The column and mobile phase were identical to those used in the Orbitrap instrument described earlier. The mass spectrometer was operated in data-dependent acquisition mode using Parallel Accumulation Serial Fragmentation (PASEF) workflow. In brief, a single full-scan mass spectrum in the time-of-flight (TOF) (350-1800 m/z, 50,000 resolution) was followed by 10 times ion release from trapped ion mobility, sequential selection by quadruple, fragmentation by HCD, and detection in the TOF (300-1500 m/z, 15,000 resolution). The total cycle time was 1.1 s, and 1/K0 for TIMS selection was set as 0.75-1.35 Vs/cm^2^. Each mass spectrum was analyzed using MSFragger for database searching against the Homo sapiens proteome database downloaded from UniProtKB (UP000005640) containing 80,581 proteins as of October 18, 2022 and label-free quantification [4-5]. Workflow LFQ-MBR was loaded for phosphoproteomics analysis with default setting of MSFragger. Fixed modifications included cysteine alkylation, and the methionine oxidation and STY phosphorylation were variable modifications. MaxLFQ was chosen for quantification approach and Min ions was set as 2. For feature detection, tolerance of m/z, retention time, and ion mobility (1/*K*_0_) were set as 10 ppm, 3 min, and 0.05, respectively. Peptides were filtered with 1% false discovery rate (FDR).

**8. LC-MS/MS for metabolomics and lipidomics**

Both metabolomic and lipidomic analyses were performed using an Agilent 1290 Infinity HPLC system coupled to an Agilent 6545 Q-tof mass spectrometer with an electrospray ion source operating in both positive and negative ion modes. For metabolomics, LC separation was carried out on a Waters BEH amide column (2.1×100 mm, 1.7 μm) at 40°C using a mobile phase consisting of MS-grade water containing 0.3% ammonium and 15 mM ammonium acetate as mobile phase A, and 90% acetonitrile with 0.3% ammonium and 15 mM ammonium acetate as mobile phase B. The flow rate was 0.3 mL/min, and the gradient of mobile phase A was changed from 10% to 50% over 8 min, then held at 50% for 2 min, changed from 50% to 10% in 0.5 min, and finally held at 10% for 6.5 min. Full scan spectra were acquired with a mass range from m/z 50 to 1200 using the high-resolution mode (Extended Dynamic Range 2GHz). The MS parameters were set as follows: capillary voltage, 3500 V in positive mode and 4000 V in negative mode; Nozzle voltage, 120 V; nebulizer gas, 35 psi; drying gas flow rate, 8 L/min; and gas temperature, 350°C. Data processing was performed using MassHunter Qualitative Workflow and Profinder 10.0 software from Agilent, and metabolite identification was conducted using accurate mass comparison and retention time matching through a home-made metabolite database.

For lipidomics, LC separation was carried out on a Waters BEH C18 column (2.1×100 mm, 1.7 μm) at 40°C using a mobile phase consisting of 10 mM ammonium acetate, 0.2 mM ammonium fluoride in 9:1 water/methanol as mobile phase A and 10 mM ammonium acetate, 0.2 mM ammonium fluoride in 2:3:5 acetonitrile/methanol/isopropanol as mobile phase B. The flow rate was 0.3 mL/min, and the gradient of mobile phase B was set to 70% for 1 min, changed from 70% to 86% over 2.5 min, held at 86% for 6.5 min, changed from 86% to 100% over 1 min, and finally held at 100% for 6 min. MS data were acquired using electrospray ionization in both positive and negative ion mode over 40-1700 m/z. MS settings were as follows: sheath gas temperature at 300℃, sheath gas flow at 11 L/min, VCap at 3500 V in positive mode and 4000 V in negative mode, Capillary at 0.09 mA, Nozzle voltage at 0 V, gas temperature at 275℃, fragmentor at 150 V, and skimmer at 65 V. Raw data were processed using Agilent Profinder 10.0 for peak detection, alignment, integration, and Lipid Annotator for annotation.

**References**

1. Smith, P.K.; Krohn, R.I.; Hermanson, G.T.; Mallia, A.K.; Gartner, F.H.; Provenzano, M.D.; Fujimoto, E.K.; Goeke, N.M.; Olson, B.J.; Klenk, D.C. Measurement of protein using bicinchoninic acid. Anal Biochem. 1985, 150, 76-85.
2. Hughes, C.S.; Moggridge, S.; Müller, T.; Sorensen, P.H.; Morin, G.B.; Krijgsveld, J. Single-pot, solid-phase-enhanced sample preparation for proteomics experiments. Nat Protoc. 2019, 14, 68-85.
3. Ding, C.; Jiang, J.; Wei, J.; Liu, W.; Zhang, W.; Liu, M.; Fu, T.; Lu, T.; Song, L.; Ying, W.; et al. A fast workflow for identification and quantification of proteomes. Mol Cell Proteomics. 2013, 12, 2370-2380.
4. Kong, A. T.; Leprevost, F. V.; Avtonomov, D. M.; Mellacheruvu, D.; Nesvizhskii, A. I. MSFragger: ultrafast and comprehensive peptide identification in mass spectrometry–based proteomics. Nat Methods. 2017, 14, 513-520.
5. Yu, F.; Haynes, S.E.; Teo, G.C.; Avtonomov, D.M.; Polasky, D.A.; Nesvizhskii, A.I. Fast Quantitative Analysis of timsTOF PASEF Data with MSFragger and IonQuant. Mol Cell Proteomics. 2020, 19, 1575-1585.

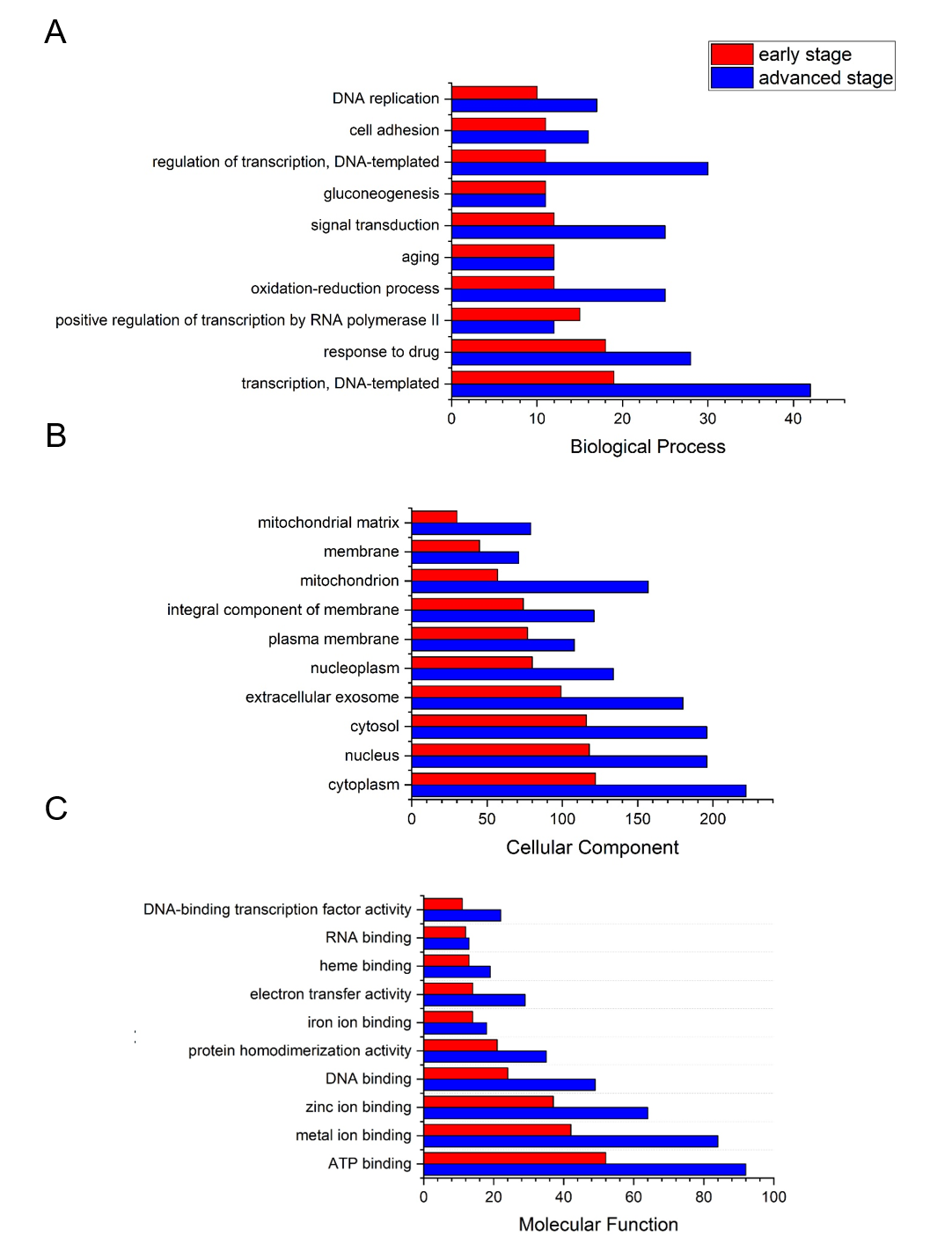

**Figure S1**. The enriched clusters using GO analysis of dysregulated proteins from proteomics. Three GO aspects of biological process, cellular component, and molecular function in early (red) and advanced (blue) stage HCC are shown in A, B, and C, respectively.

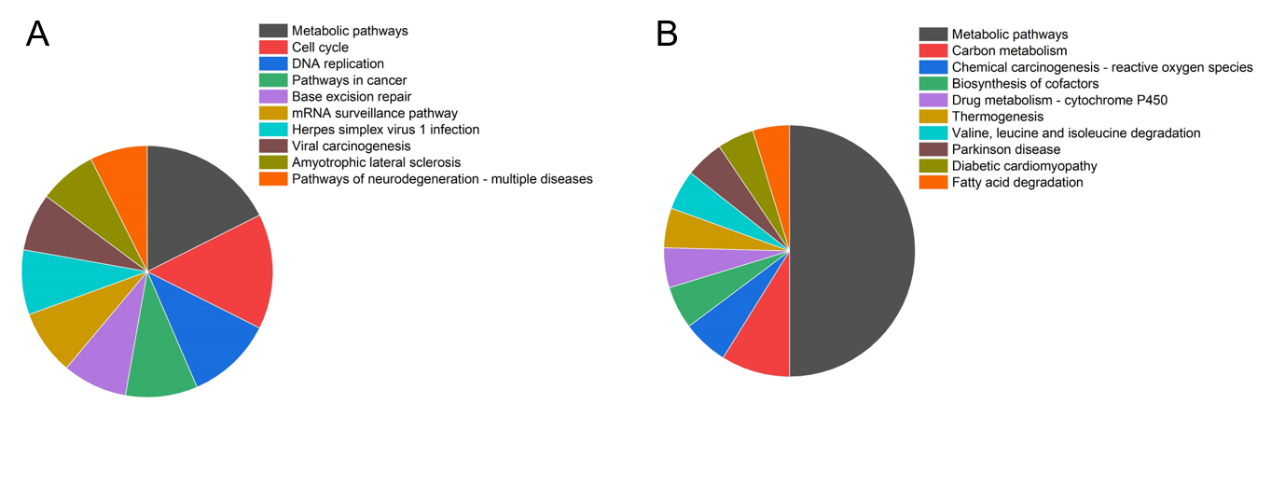

**Figure S2**. The top 10 KEGG enriched pathways in up-regulated proteins (A) and down-regulated proteins (B).

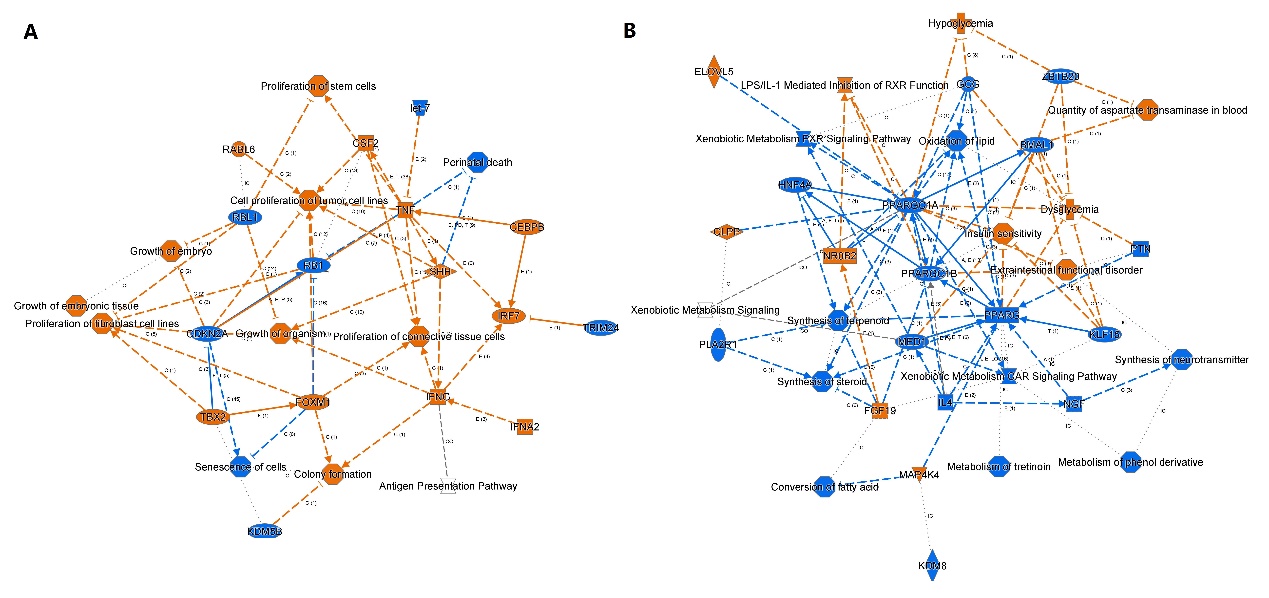

**Figure S3**. The enriched bioprocesses for up-regulated (A) and down-regulated (B) molecules in early-stage HCC by PML assay using IPA.

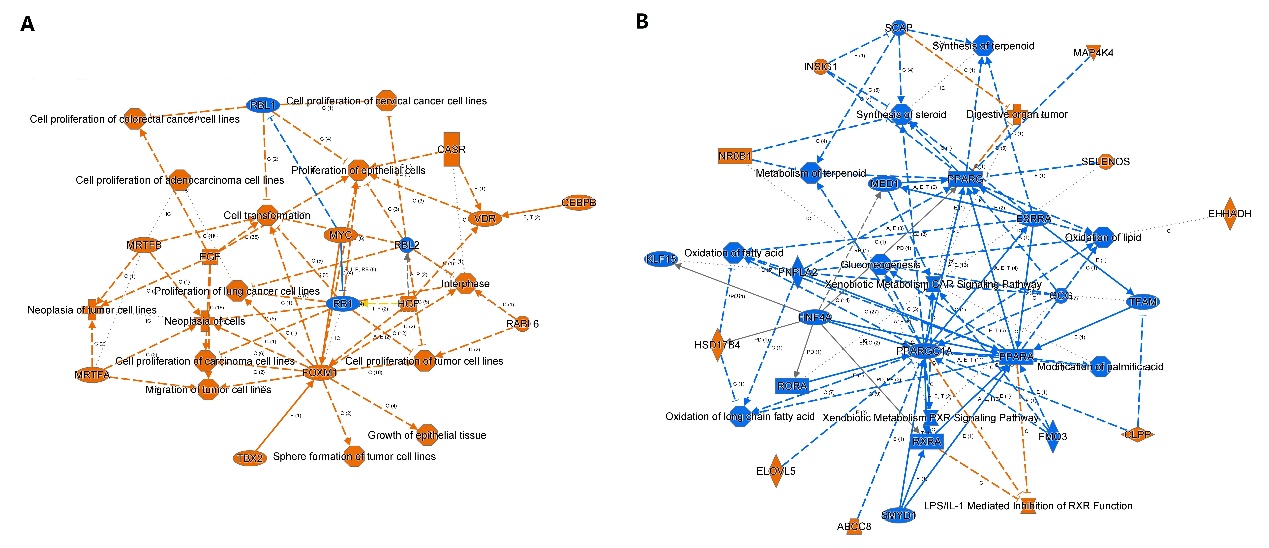

**Figure S4**. Enriched bioprocesses for up-regulated (A) and down-regulated (B) molecules in advanced-stage HCC by PML assay using IPA.

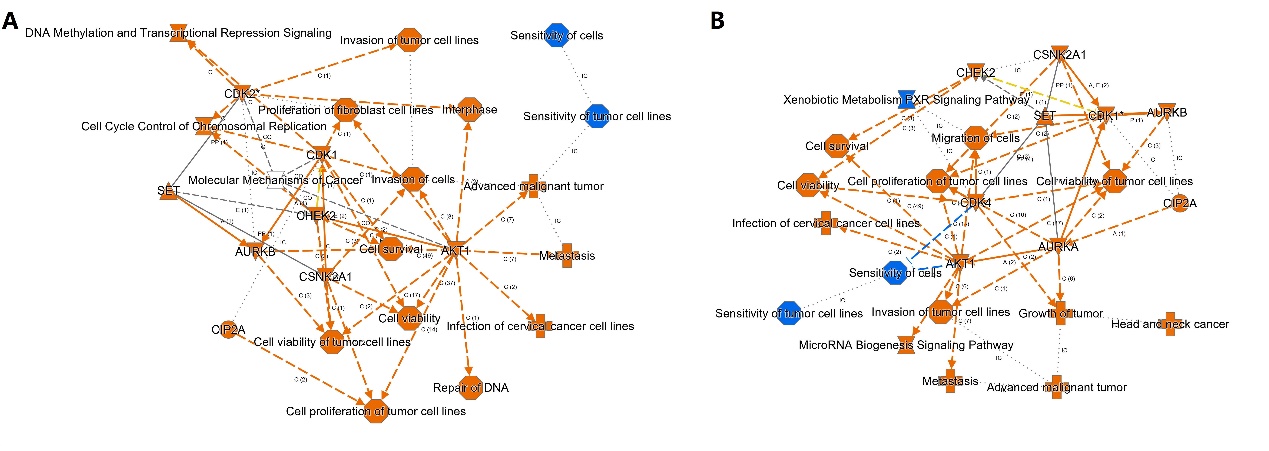

**Figure S5**. The enriched bioprocesses for dysregulated phosphopeptides in early (A) and advanced (B) stage HCC by phosphorylation assay using IPA.

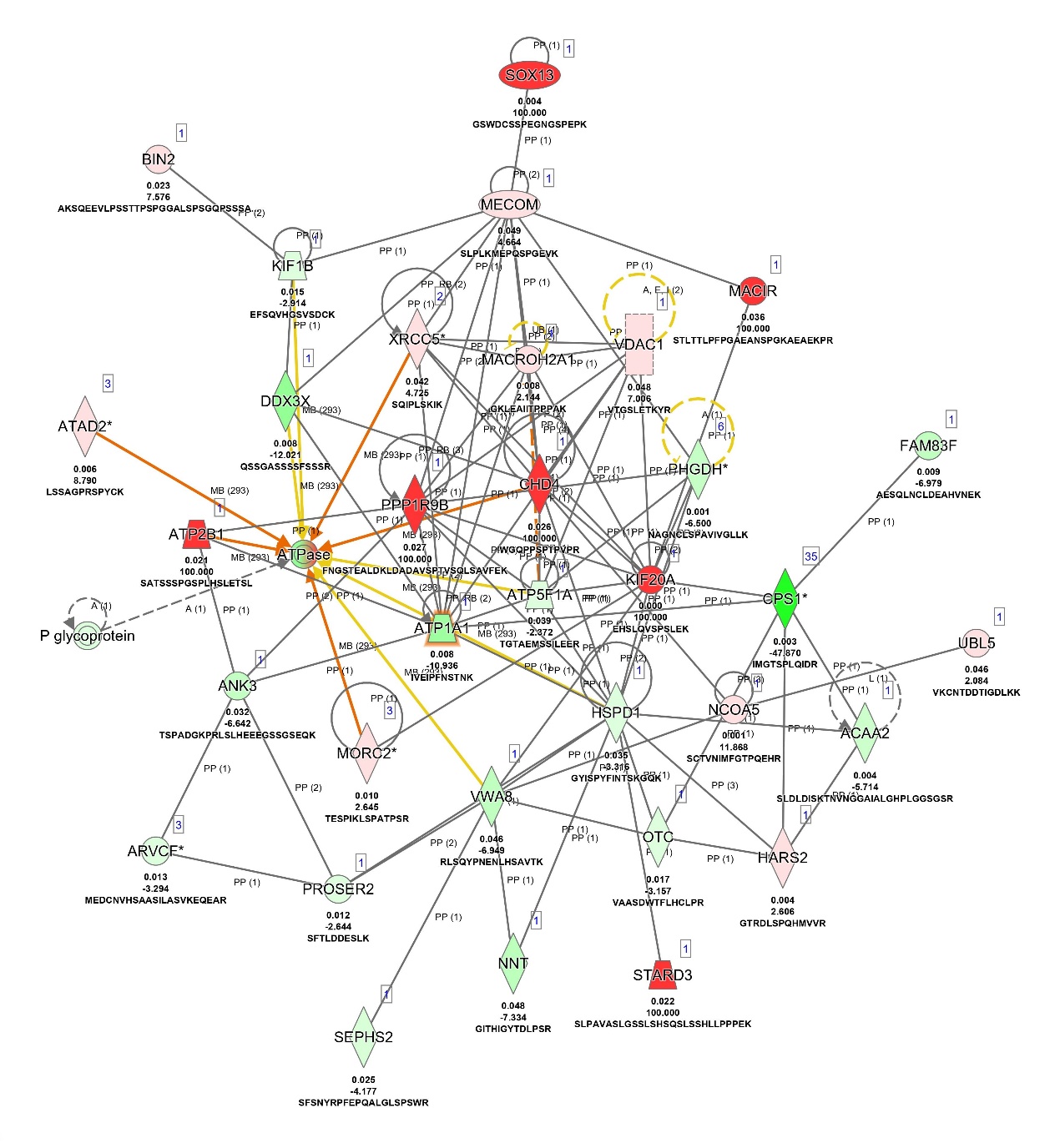

**Figure S6**. An example of protein phosphorylation interaction network based on IPA analysis.

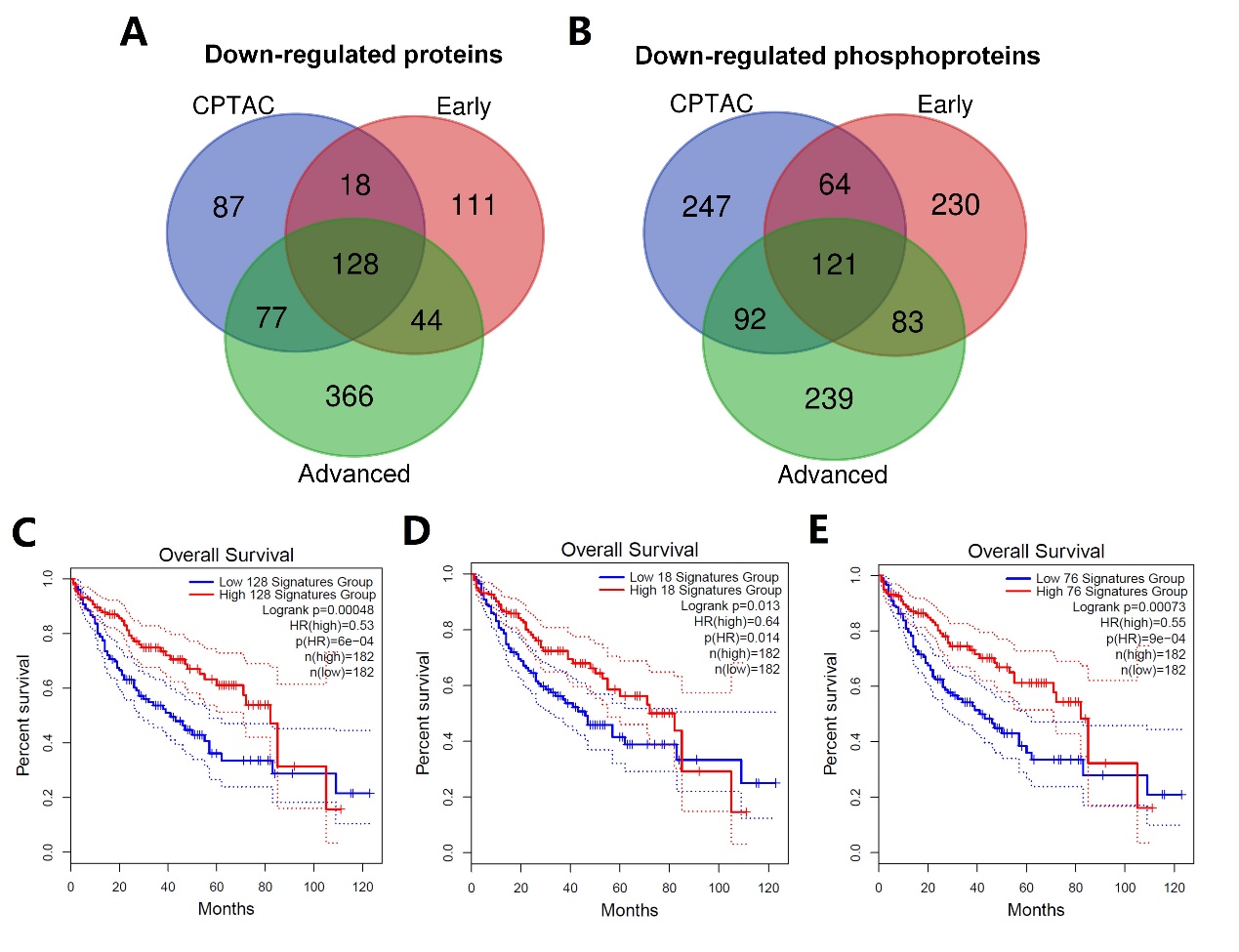

**Figure S7**. The comparisons with CPTAC data for down-regulation. Venn diagram showing the overlap between the down-regulated proteins (A) and phosphoproteins (B) analyzed from CPTAC database, early- and advanced stage HCC data in current study. Kaplan–Meier analysis for the overall survival of the 128- (C), 18- (D) and 76- (E) gene signatures.
